# Cellular, Circuit and Transcriptional Framework for Modulation of Itch in the Central Amygdala

**DOI:** 10.1101/2021.02.18.431884

**Authors:** Vijay K Samineni, Jose G. Grajales-Reyes, Gary E Grajales-Reyes, Eric Tycksen, Bryan A Copits, Christian Pedersen, Edem S Ankudey, Julian N Sackey, Sienna B Sewell, Michael R Bruchas, Robert W. Gereau

## Abstract

Itch is an unpleasant sensation that elicits robust scratching and active avoidance. However, the identity of the cells and neural circuits that organize this information remains elusive. Here we show the necessity and sufficiency of itch-activated neurons in the central amygdala (CeA) for both itch sensation and active avoidance. Further, we show that itch-activated CeA neurons play important roles in itch-related comorbidities, including anxiety-like behaviors, but not in some aversive and appetitive behaviors previously ascribed to CeA neurons. RNA-sequencing of itch-activated CeA neurons identified several differentially expressed genes as well as potential key signaling pathways in regulating pruritis. Finally, viral tracing experiments demonstrate that these neurons send a critical projection to the periaqueductal gray to mediate modulation of itch. These findings reveal a cellular and circuit signature of CeA neurons orchestrating behavioral and affective responses to pruritus in mice.

## Main

As organisms have evolved, it has been essential that they acquire the means to sense physical and chemical threats in the world around them. One such threat detection system is itch, which accompanies unpleasant sensations that evoke strong urges to scratch and promote active avoidance behavior (Bautista et al., 2014; Han and Dong, 2014; Ikoma et al., 2006; LaMotte et al., 2014). Orchestrating adaptive behaviors (e.g. scratching an itch, avoidance of active threats) in the future requires rapid routing of information to brain regions that can encode memories and modify behavior based on prior experiences. The central amygdala (CeA) represents a strong candidate for these functions, as the CeA is thought to play a critical role in learning and modifying sensory and emotional memories and translating this information into apt adaptive behaviors (Fadok et al., 2018; Grundemann and Luthi, 2015; LeDoux and Daw, 2018). Recent studies have implicated the CeA in the regulation of itch (Albisetti et al., 2019; Chen et al., 2016; Ehling et al., 2018; Mu et al., 2017), and elevated activity in the CeA has been seen in patients during experimental itch (Papoiu et al., 2014; Vierow et al., 2015). Nevertheless, it is currently unknown how CeA neurons encode and modify the sensory or emotional components of itch. To address these questions, we used optical imaging, activity dependent labelling, neural tracing and cell activity-specific RNA sequencing to systematically investigate itch-activated CeA neurons and their projections in eliciting itch and its related comorbidities.

## Results

To assess the potential activation of CeA neurons by itch in real time, we performed fiber photometry recordings to follow CeA neural activity during pruritogen-evoked scratching in awake, behaving mice. To record real-time Ca^2+^ dynamics in the CeA (Cui et al., 2013), we expressed the genetically-encoded calcium indicator, GCaMP6s, in CeA GABAergic neurons, using viral delivery of Cre-dependent GCaMP6s in Vgat-IRES-Cre mice (Swanson and Petrovich, 1998) (Fig. 1a and Supplementary Fig. 1a-c). Subcutaneous injection of chloroquine in the nape of the neck induced scratching behavior and resulted in robust increases in CeA neuronal activity (Fig. 1b). This activity commenced with initiation of scratching and stabilized whenever scratching stopped (Figure 1c-e), suggesting that the elevated activity was tightly-coupled with the act of scratching.

**Fig 1.**
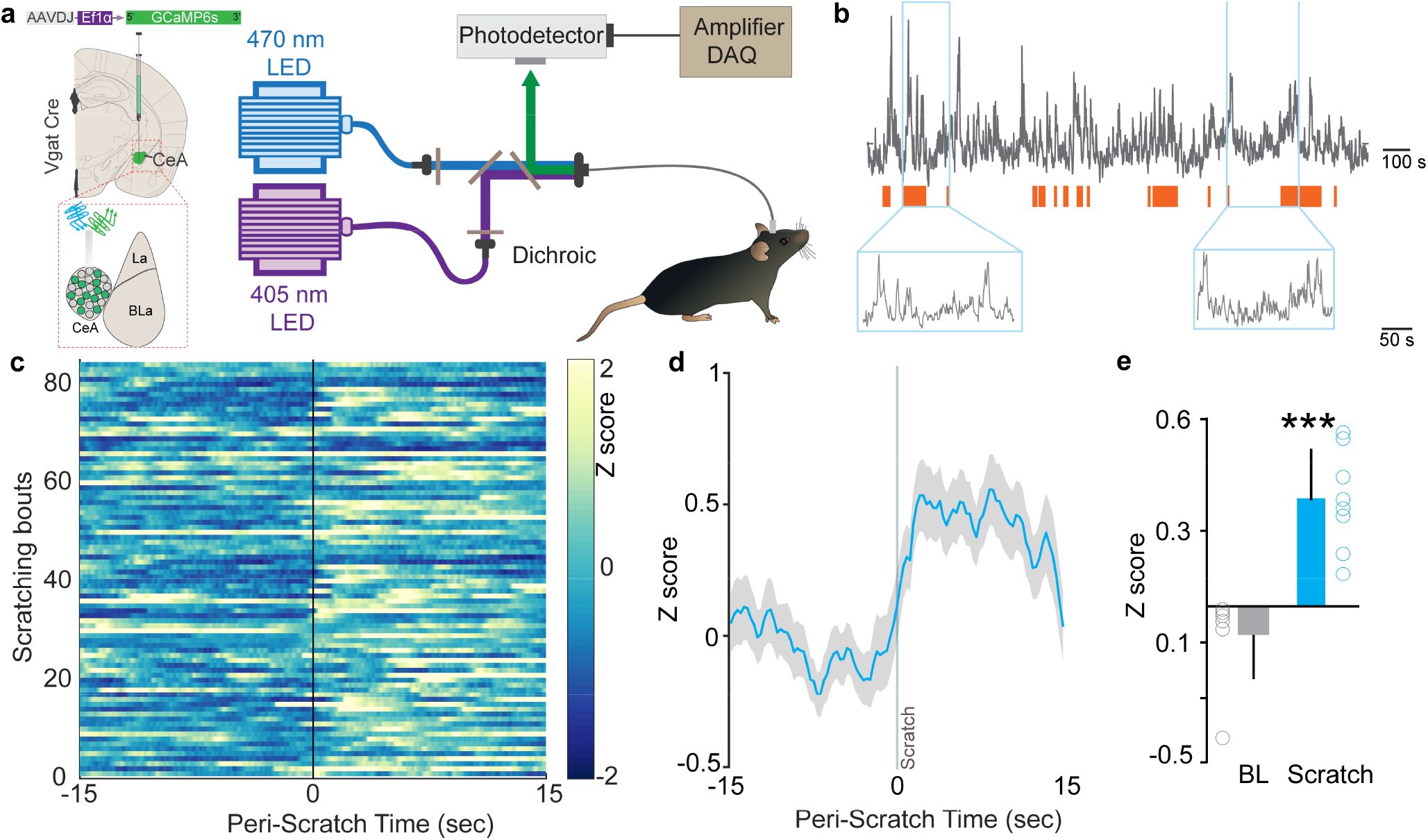
Neural dynamics of itch activated with CeA neurons. (a) Scheme demonstrating viral injection strategy and fiber placement to record CeAVgat neural activity in response to chloroquine. (b) Raw Ca2+ dynamics recorded from CeAVgat neurons and their relationship to chloroquine evoked scratching bouts (orange bars). (c) Heatmap showing Ca2+ dynamics of all trials of Vgat+ve vlPAG neurons relative to the initiation of chloroquine evoked scratching bouts (time zero). (d) Averaged GCaMP6s fluorescence signal of CeAVgat neurons showing rapid increases in fluorescence on the initiation of scratching bouts. Trace plotted as mean (blue line) ± SEM (gray shading), and the vertical line indicates initiation of scratching bouts. (e) Chloroquine-evoked scratching resulted in a significant increase in CeAVgat neuronal activity as measured by this change in GCaMP6s fluorescence. **(N=**8, t test, t=5.923, df=14, P <0.0001).

Consistent with these real-time dynamic recordings, activity-dependent mapping studies show robust cFos labelling bilaterally in the CeA following chloroquine injection in the nape of the neck compared to saline-injected mice (Supplementary Fig. 1a, b). We observed no significant differences in cFos labeling between right and left CeA (Supplementary Fig. 1c).

These observations provide cellular confirmation of prior reports (Mochizuki et al., 2014; Mochizuki et al., 2003; Papoiu et al., 2013) that indicated a possible role for the CeA in itch processing, but the underlying neural circuitry remains to be identified. If CeA neurons function as a key node in the circuit that tightly regulates sensory and affective component of itch, then their activation should trigger potentiation of the itch-scratching cycle and its aversive state. CeA neurons are molecularly heterogeneous and mediate diverse behaviors generally related to negative affect (John et al., 2015; Kalin et al., 2004; LeDoux, 2003; Ressler and Mayberg, 2007; Roozendaal et al., 2009; Tye et al., 2011), so we reasoned global manipulation of CeA neuronal activity would not provide the specificity needed to test the specific roles of itch-activated neurons. To enable the desired selective manipulation of itch-specific neuronal populations in the CeA, we used “Targeted Recombination in Active Populations” mice (Guenthner et al., 2013). These mice express the tamoxifen-dependent CreER^T2^ recombinase from the *Fos* promoter. CreER^T2^ expression is induced in neurons that were recently active. Catalytic activity of CreER^T2^ is stabilized in the presence of 4-hydroxytamoxifen (4-OHT), resulting in transgene recombination. By timing the administration of 4-OHT to coincide with recently increased neuronal activity during acute chloroquine stimuli, we can gain permanent genetic access to itch-responsive CeA neurons (aka FosTRAP mice). To test the validity of this approach, we crossed FosTRAP mice to a Cre-dependent tdTomato flox-stop reporter line (Madisen et al., 2010). We injected chloroquine or saline into the nape of the neck, paired with injection of 4-hydroxytamoxifen (4-OHT) to induce Cre-mediated recombination of the tdTomato in activated (cFos-expressing) neurons (Fig. 2a & b). FosTRAPing with chloroquine treatment produced robust tdTomato expression in both the right and left CeA (Fig. 2c, d & e), but not saline-treated controls (Supplementary Fig. 2e-g), consistent with our cFos staining results above. To confirm that the FosTRAPed neurons are specific to the chloroquine-evoked scratching, one-week post-FosTRAP, we immunostained for c-Fos protein in mice that received an additional chloroquine injection just prior to sacrificing (Fig. 2c & f). The majority of the tdTomato-positive CeA FosTRAPed neurons faithfully overlap with cFos-positive cells. These results demonstrate that we can efficiently gain genetic access to neurons that are activated by chloroquine.

**Fig 2.**
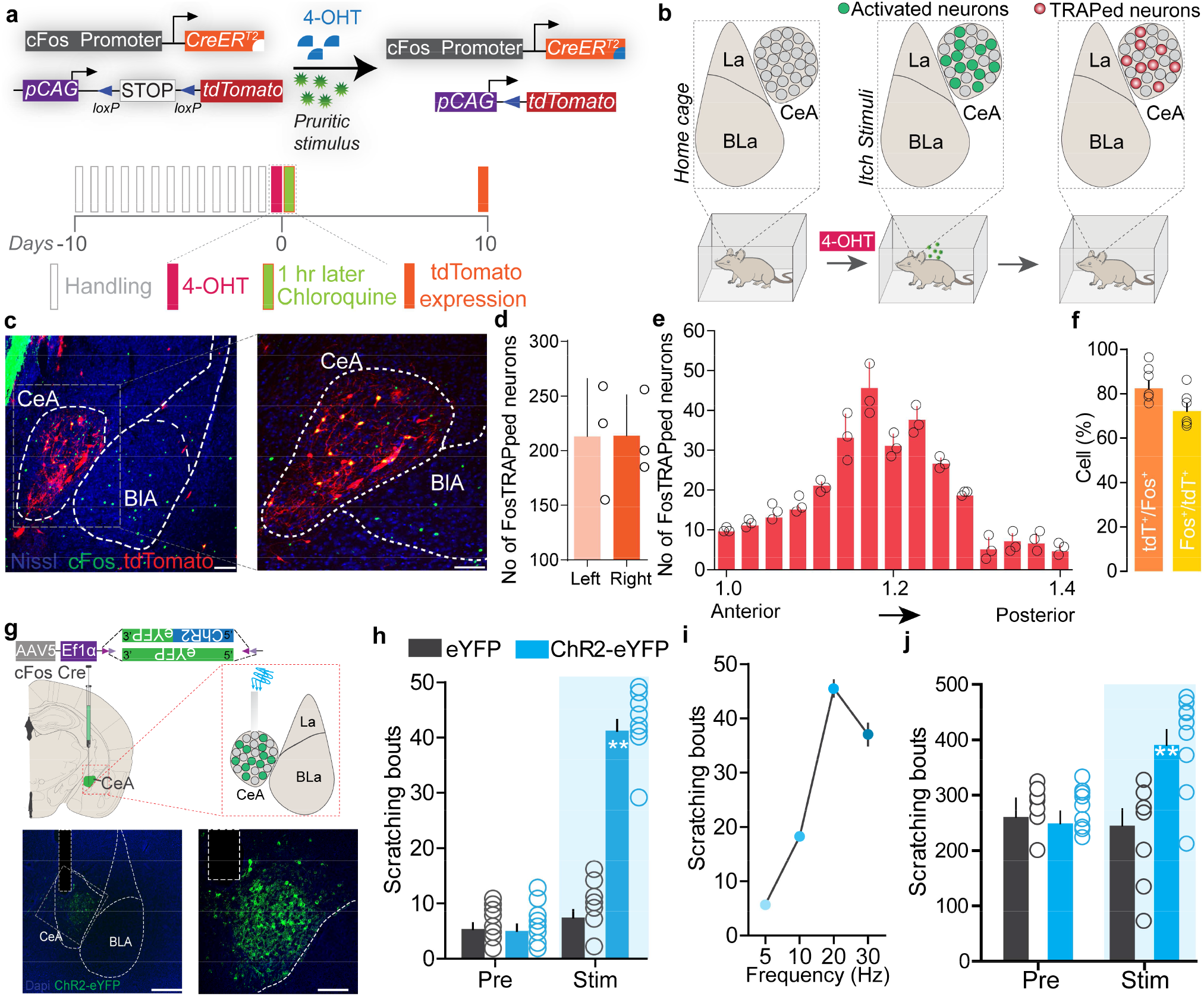
Itch-activated CeA neurons can drive pruritic behaviors. (a) FosTRAP strategy to selectively label itch-activated neurons in the CeA. (b) Scheme illustrating experimental strategy. (c) FosTRAPping with chloroquine-evoked scratching produces robust tdTomato expression in the CeA. Colocalization of itch-TRAPed neurons (red) in the CeA with cFos immunoreactivity (green) following a 2nd administration of chloroquine 7 days post. Scale bar= 85 and 250 μm. (d) Quantification of the number of FosTRAPped neurons in left and right CeA after chloroquine injection. n = 3 per group. t test, t=0.4339, df=2, p =0.70. (e) Rostro-caudal distribution of itch-TRAPed CeA neurons after chloroquine injection_ (f) Colocalization of chloroquine-activated cFos with tdTomato+ve itch-TRAPed neurons. Relative percentages of Fos+ve neurons that are tdTomato+ve and tdTomato+ve neurons that are Fos+ve. n = 6 per group. t test, t=2.04 df=10, p =0.048. (g) Scheme to selectively express optogenetic constructs in itch-TRAPed CeA neurons_ Illustration and representative section showing fiber optic placement above FosTRAPped CeA neurons expressing ChR2-eYFP (green). Scale bar, 100 μm. (h) Photostimulation (20 Hz) of itch-TRAPed CeA neurons produces robust spontaneous scratching. n = 6-11 per group. Pre Vs. Stirn, F (1,30) = 3; eYFP Vs. ChR2, F (1, 14) = 3.24, p = 0.0001. (i) Increases in scratching are frequency dependent. n = 6 per group. U) Optical activation of itch-TRAPed CeA neurons potentiates chloroquine-evoked scratching while no changes were observed in control mice. n = 7 per group. Pre Vs. Stirn, F (1,12) = 33.15; BL Vs. Stirn in ChR2, F **(1,**6) = 6.915, p = P=0.0391.

To test whether reactivating itch-responsive CeA neurons can recapitulate itch behaviors, we expressed the Cre-dependent excitatory opsin, ChR2 (AAV5-EF1a-DIO-ChR2-eYFP) or a control virus (AAV5-EF1a-DIO-eYFP) in the right CeA of FosTRAP mice and FosTRAPed with chloroquine treatment as above (Fig. 2g). This produces expression of ChR2 specifically in CeA neurons responsive to itch, enabling their selective light-dependent activation. Optogenetic reactivation of FosTRAPed (ChR2^+^) right CeA neurons resulted in significant spontaneous scratching and grooming behaviors compared to pre-stimulation baseline and photostimulation of eYFP-expressing control mice. Interestingly, although ChR2 was FosTRAPed by injecting chloroquine into the nape of the neck, we observed spontaneous scratching and grooming behaviors directed all over the body (data not shown) in a stimulation frequency-dependent manner (Fig. 2h, i). Even though some functions of the CeA are lateralized (Carrasquillo and Gereau, 2007), elicitation of itch behaviors are not lateralized to the right CeA as optical stimulation of FosTRAPed ChR2^+^ neurons in the left CeA also resulted in significant spontaneous scratching behaviors compared to pre-stimulation baseline and photostimulation of eYFP-expressing controls (Supplementary Fig. 3). This result is consistent with the observation that chloroquine injection induces cFos expression in left and right CeA (Supplementary Fig. 2d). To further confirm these results and as a complementary approach, we expressed the Cre-dependent excitatory DREADD, hM3Dq (AAV5-hSyn-DIO-hM3Dq-mCh) or a control virus (AAV5-hSyn-DIO-mCh) in the CeA of FosTRAP mice. Chemogenetic activation of FosTRAPed CeA neurons also resulted in significant spontaneous scratching behaviors (Supplementary Fig. 4e), consistent with the optogenetic results.

In contrast, stimulation of FosTRAPed neurons has no significant effect on hindpaw thermal sensitivity (Supplementary Fig. 5b) or licking and biting behaviors Supplementary Fig. 4f). However, stimulation of FosTRAPed neurons slightly increased mechanical sensitivity suggesting that these neurons can encode generalized itch-related behavior and hypersensitivity to mechanical stimuli (Supplementary Fig. 5c). These results suggest that FosTRAPed neurons might be involved in nociceptive processing (Neugebauer and Li, 2002). To further determine how reactivation of itch-activated (ChR2^+^) FosTRAPed neurons (hereafter referred to as “itch-TRAPed neurons”) can affect ongoing itch behaviors, we administered chloroquine and optically activated itch-TRAPed CeA neurons. Chloroquine evoked scratching was potentiated with optical reactivation of CeA itch-TRAPed neurons while no changes were observed in the eYFP controls (Fig. 2j). Chemogenetic stimulation of itch-TRAPed neurons produced similar results (Supplementary Fig. 4h).

Itch is an aversive sensory experience in humans and rodents (Desbordes et al., 2015; Mochizuki et al., 2015; Mochizuki et al., 2014; Papoiu et al., 2012; Papoiu et al., 2013), and the CeA mediates aversive phenotypes (Carrasquillo and Gereau, 2007; Ciocchi et al., 2010; Ehrlich et al., 2009; Haubensak et al., 2010; Tovote et al., 2016). Therefore, we wanted to assess whether itch-TRAPed CeA neurons encode negative valence associated with itch. We performed closed-loop real-time place-testing (RTPT) to assess affective state, where an animal freely explores two chambers but receives photostimulation of ChR2^+ve^ itch-TRAPed neurons in only one chamber. Reactivation of itch-TRAPed neurons produced robust place aversion to the stimulated side of the chamber while eYFP-FosTRAPed controls did not (Fig. 3b-d), thus demonstrating that itch-activated CeA neurons carry negative reinforcement signals.

**Fig. 3.**
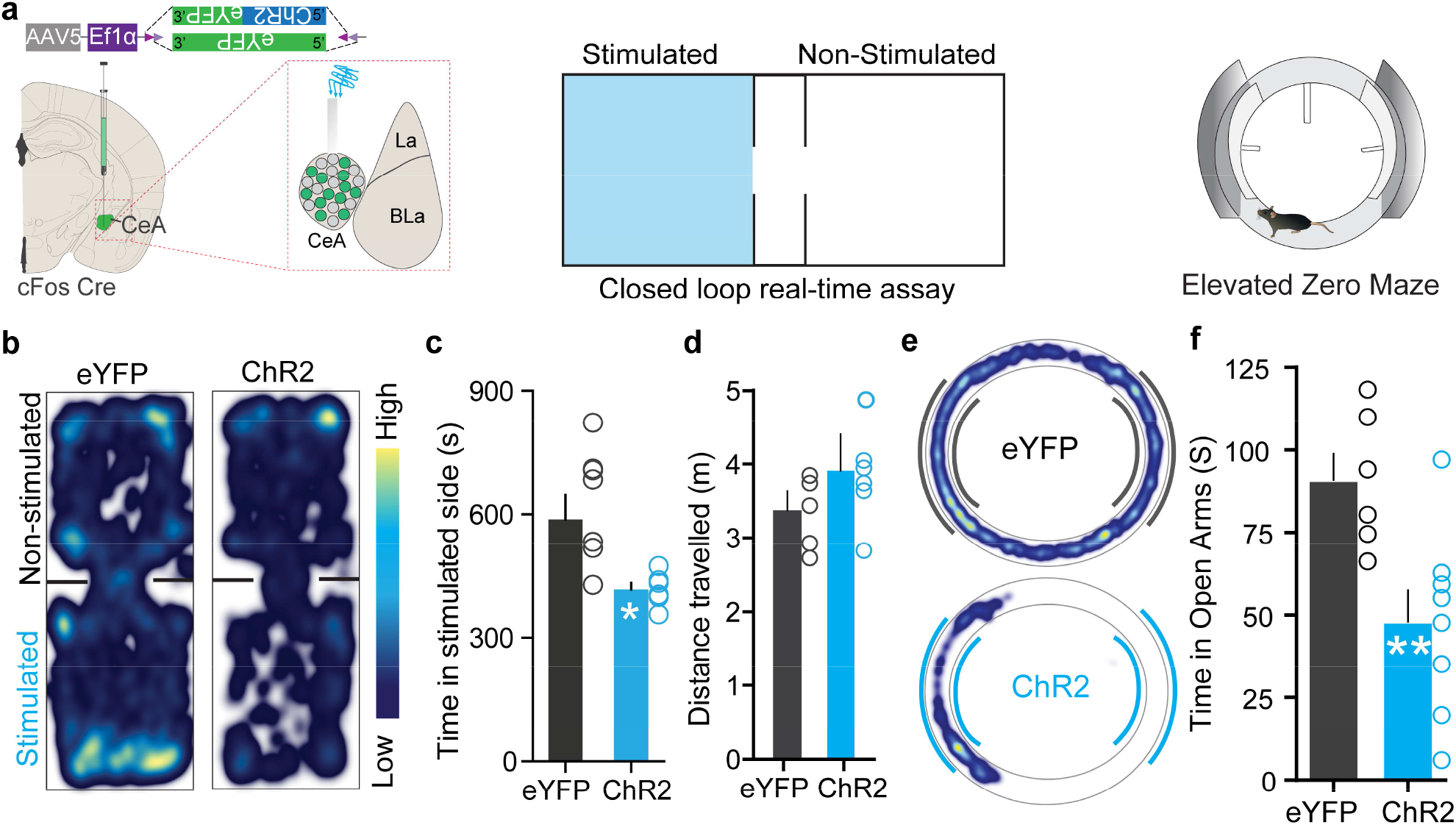
Itch-activated CeA neurons are negatively reinforcing. (a) Illustration of strategy to express ChR2/eYFP selectively in itch-TRAPed neurons of the CeA. Experimental schematic of closed loop real-time assay and elevated zero maze. (b) Real-time place aversion assay with spatial location heatmaps of ChR2 and eYFP mice during closed loop optical stimulation. t test, t=2.806, df=12, p =0.0159. (c) Total time spent and (d) distance travelled in the photostimulation-paired chamber for ChR2 and eYFP mice. n = 7 per group, t test, t=0.7510, df=12, p =0.4142. (e) Representative occupancy heat map showing spatial location in the elevated zero maze (EZM)of a control mouse (eYFP) and a mouse injected with DIO-ChR2. (f) Optogenetic activation of itch-TRAPed CeA neurons causes a significant reduction in time spent in open arms in EZM. Light stimulation was delivered entire time mice were on EZM n= 6-10 per group. t test, t=5.922, df=12, p =0.0086.

Patients with pruritic skin disorders exhibit heightened anxiety (Ginsburg, 1995) and prior studies have shown CeA as a critical hub in coordinating anxiety states (Ahrens et al., 2018; Shackman and Fox, 2016). Therefore, we evaluated if reactivation of itch-TRAPed CeA neurons can drive anxiety-like behavior using the elevated zero maze (EZM) assay and open-field test (OFT). Optogenetic and chemogenetic reactivation of itch-TRAPed neurons lead to a profound decrease in time spent in the open arms of EZM compared to controls, indicating anxiogenic-like behavioral state (Fig. 3e-, Supplementary Fig. 6b, c). Reactivation of these neurons also lead to decreased time spent in the center during the OFT, further suggesting these neurons can drive anxiety-like behavior (Supplementary Fig. 6d-i). Notably, opto- and chemogenetic reactivation of FosTRAPed neurons did not drive freezing or flight responses in open-field test (Supplementary Fig. 6e & g), suggesting these neurons are not involved in fear-like behaviors. Furthermore, stimulation of these neurons also had no effect on feeding and other appetitive behaviors the CeA is reported to evoke (Douglass et al., 2017; Han et al., 2017; Kim et al., 2017; Li et al., 2013 (Supplementary Fig. 7a-c & Supplementary Fig. 8).

Having shown that itch-TRAPed CeA neurons are sufficient to drive itch-related sensory and affective behaviors, we aimed to determine if endogenous activity of these neurons is necessary for itch-related behaviors. We selectively inhibited itch-responsive CeA neurons by expressing the Cre-dependent inhibitory DREADD, hM4Di (AAV5-hSyn-DIO-hM4Di-mCh), or a control virus (AAV5-hSyn-DIO-mCh) in itch-activated CeA neurons of FosTRAP mice (Fig. 4a-c). CNO application to ex vivo CeA slices from itch-TRAPed mice decreased neuronal excitability to supratheshold stimuli (Fig. 4d). Chemogenetic inhibition of CeA itch-TRAPed neurons by CNO injection significantly attenuated chloroquine-evoked scratching compared to pre-CNO baseline and compared to mCherry-expressing controls (Fig. 4e, f). These results suggest that itch-activated (itch-TRAPed) CeA neurons are necessary for chloroquine induced scratching behavior. We observed no significant effect on thermal or mechanical sensitivity by inhibiting CeA itch-TRAPed neurons (Supplementary Fig. 5d, e). Furthermore, silencing CeA itch-TRAPed neurons did not lead to freezing or flight responses or anxiolytic effects (Supplementary Fig. 6j-n). Inhibition of itch-TRAPed neurons also did not affect feeding (Supplementary Fig. 7d) and appetitive behaviors (Supplementary Fig. 8). Because activating CeA itch-TRAPed neurons lead to robust avoidance, we hypothesized that inhibiting these neurons would block conditioned place aversion (CPA) to chloroquine. We performed CPA to chloroquine in itch-TRAPed mice expressing either hM4Di or mCherry in itch-responsive neurons (Fig. 4g). Silencing itch-TRAPed (hM4Di^+ve^) CeA neurons with CNO injection in the chloroquine-paired chamber during conditioning blocked CPA to chloroquine, while CNO-treated mCherry controls exhibited CPA to chloroquine, suggesting that these neurons can robustly modulate the aversive component of itch (Fig. 4h-k).

**Fig 4.**
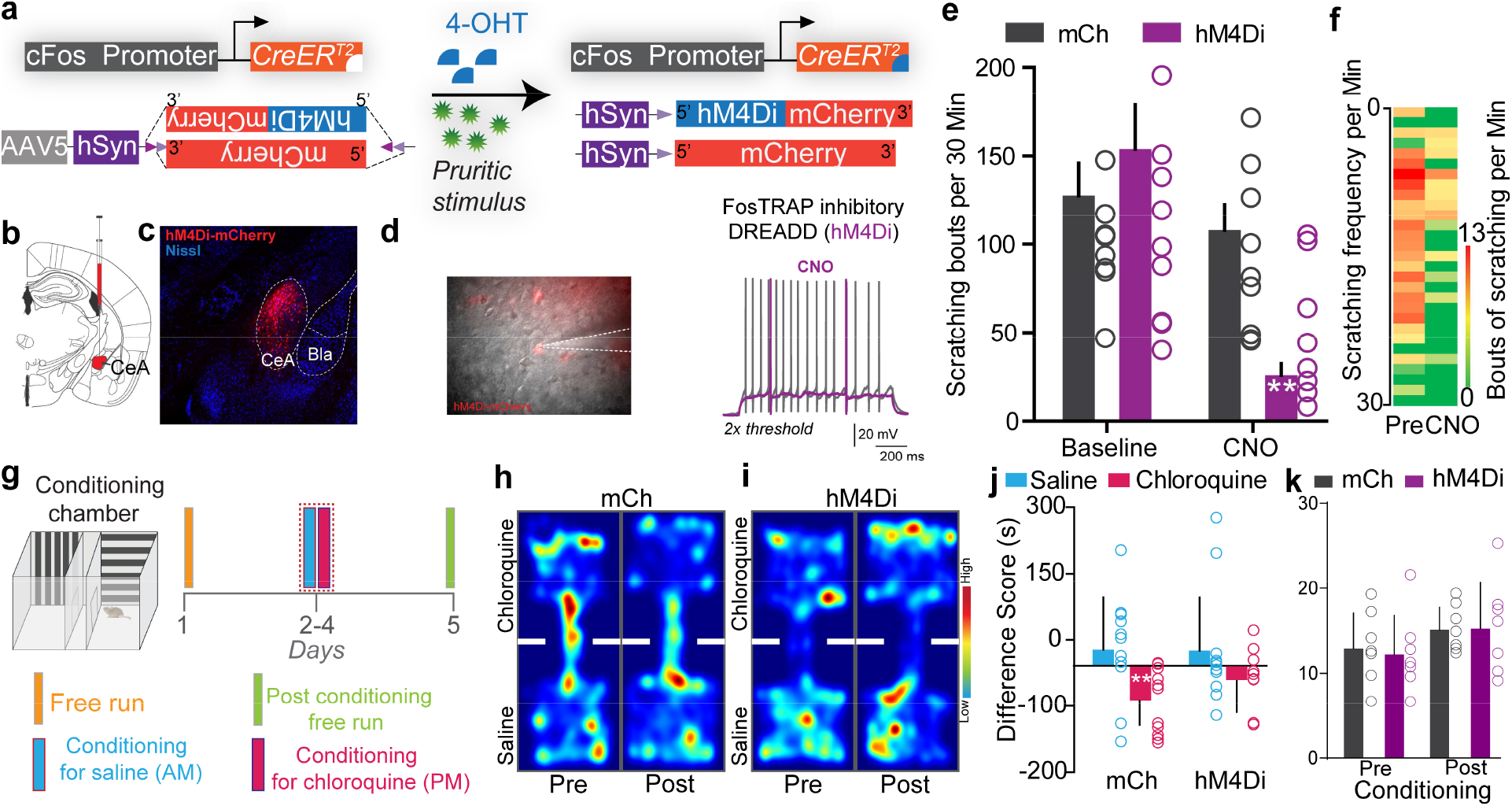
Inhibiting itch-activated CeA neurons impairs aversive learning associated with itch. (a) Illustration of strategy to express inhibitory DREADDs selectively in itch-TRAPed neurons of the CeA. (b) Experimental timeline to FosTRAP DREADDs in CeA neurons. (c) Representative section showing itch-TRAPed CeA neurons expressing hM4Di-mCherry (Red). Scale bar, 75 μm. (d) Infrared DIC image of CeA itch-TRAPed neurons expressing hM4Di-mCherry. In hM4Di+ve CeA neurons, CNO bath application decreased neuronal excitability to supratheshold stimuli. (e) Chemogenetic inhibition of itch-TRAPed CeA neurons leads to a significant reduction in chloroquine evoked scratching. CNO has no effect on chloroquine-evoked scratching in control mice expressing mCherry. n = 8-9 per group. p = 0.0011. (f) Heat map showing averaged chloroquine-evoked scratching bouts pre- and post-CNO in mice expressing hM4Di in CeA. (g) Schematic and timeline of conditioned place aversion experimental design with chemogenetic silencing. Representative heat map showing spatial location of a control mouse injected with the DIO-mCh (h) and the DIO-hM4Di DREADD virus (i), pre- and post-chloroquine conditioning. U) Change in chamber occupancy time in the chloroquine-paired chamber compared to the saline-paired chamber after chemogenetic silencing. n = 11 per group. p = 0.044. (K) Distance travelled in chloroquine paired chamber did not differ pre- and post-conditioning in mCh and hM4Di mice. n = 0.769 per group.

We next sought to understand the circuit context of these itch-activated CeA neurons and explored the downstream nodes that might mediate expression of scratching behaviors. We found that itch-TRAPed CeA neurons send notably dense axonal projections in the ventral periaqueductal gray (vPAG) (Fig. 5a-d). We confirmed these results by injection CTB into the vPAG (Supplementary Fig. 9a, b) and also by injecting retro Cre DIO GFP in to the vPAG in Vgat cre mice (Supplementary Fig. 9c, d). Injection of RV-GFP into vlPAG of Vgat and Vglut2 Cre mice labeled monosynaptic projections from the CeA consistent with prior work (Supplementary Fig. 9e-k). Because the vPAG has previously been shown to contribute to pruritic behaviors (Gao et al., 2019; Samineni et al., 2019), we focused our functional studies on this CeA→vPAG circuit. If this CeA→vPAG circuit mediates scratching behaviors elicited by the itch-TRAPed CeA neurons, then stimulating this projection should recapitulate these behaviors. As predicted, photostimulating ChR2-expressing itch-TRAPed CeA neuronal terminals in the vPAG recapitulated spontaneous scratching behaviors (Fig. 5e). Activating itch-TRAPed CeA neuronal terminals in the vPAG did not produce freezing or flight responses. To determine if reactivation of itch-TRAPed CeA→vPAG projections (ChR2^+^) can influence ongoing itch behaviors, we administered chloroquine and optically activated itch-TRAPed CeA→vPAG projections. Chloroquine evoked scratching was potentiated with optical reactivation of CeA→vPAG projections while no changes were observed in the eYFP controls (Fig. 5f). These results show that the CeA→vPAG neuronal circuit is crucial node in mediating pruritic behaviors.

**Fig 5.**
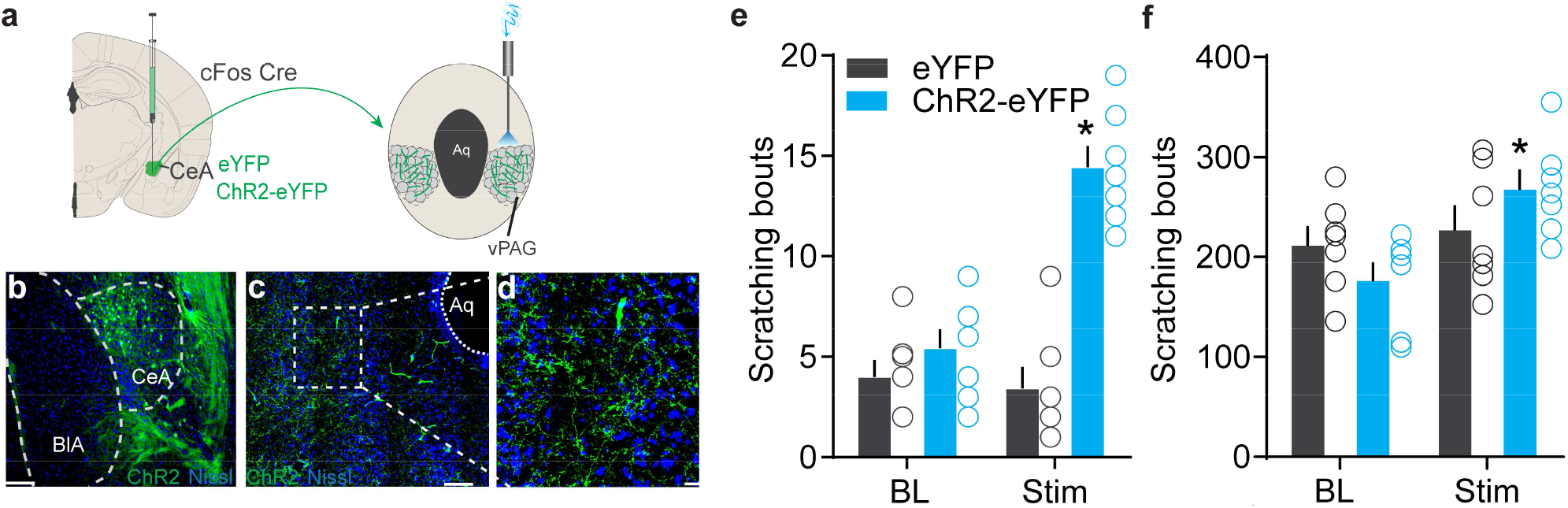
Identification of the downstream circuit of itch-activated CeA neurons. (a) Scheme showing expression of ChR2 in itch-TRAPed CeA neurons and their axonal photostimulation in the vPAG. (b) FosTRAPped CeA neurons expressing ChR2-eYFP. Scale bar, 125 μm. (c, d) ltch-TRAPed ChR2+ve CeA axonal terminals ramify densely in the vPAG. Scale bar, 100 and 25 μm. (e) Optogenetic stimulation of FosTRAPed ChR2+ve axonal projections from CeA in the vPAG resulted in significant spontaneous scratching, whereas photostimulation had no effect on scratching in control mice. Pre vs Stirn, F (1, 12) = 33.15, p <0.0001, n = 5-7 per group. (f) Optical activation of itch-TRAPed CeA neurons potentiates chloroquine-evoked scratching while no changes were observed in control mice. n = 7 per group. BL Vs. Stirn in eYFP, F (1,6) = 0.019, P=0.8924; BL Vs. Stirn in ChR2, F (1, 6) = 9.109, P=0.0235.

Lastly, we performed RNA-seq to identify transcriptional profiles of itch-activated CeA cells (Fig. 6a). To do this, we TRAPed tdTomato in itch-activated CeA neurons as described above, and separated the itch-TRAPed neurons from adjacent tdTomato^-ve^ cells for comparative RNA-seq analysis. Correlation analysis of RNA-seq data revealed itch-TRAPed tdTomato^+ve^ cells and TRAPed tdTomato^-ve^ cells are clustered apart from each other (Supplementary Fig. 11c). In our sequencing results, we observed that both the tdTomato^+ve^ and tdTomato^-ve^ cells expressed Slc32A1 transcript (VGAT, a marker for GABAergic neurons), consistent with the notion that the majority of CeA neurons are GABAergic. Hierarchical clustering analysis of genes shows highly correlated gene expression patterns that show unique expression profiles in FosTRAPed^+ve^ CeA neurons vs FosTRAPed^-ve^ CeA neurons (Supplementary Fig. 11d). We identified numerous highly correlated gene clusters based on their expression levels in FosTRAPed^+ve^ neurons (Supplementary Fig. 11e). Subsequent analysis of itch-TRAPed neurons revealed significant enrichment of several unique transcripts in the itch-activated neurons (Fig. 6b). Weighted gene correlation network analysis (WGCNA) of genes identified a cluster of up-regulated genes 99% correlated and highly significant for itch-activated neurons (Supplementary Fig. 11f-h). To link transcriptional profiles of FosTRAPed cells to known CeA functional pathways, we performed pathway analysis. From KEGG and Gene Set Enrichment Analysis (GSEA), we have identified changes in the expression of functionally related candidate genes that are enriched in several pathways (Fig. 6c). We have identified significantly enriched CeA candidate genes that might be associated with pruritus regulation, as well as significantly genes expressed at significantly lower levels relative to the non-TRAPped cells that could be involved in the suppression of pruritus. To independently confirm our RNAseq findings, we performed dual-color fluorescent in situ hybridization (FISH) to visualize mRNA expression of several candidate genes enriched specifically only in the fos-positive cells induced by pruritic stimuli. We observed considerable overlap between NTSR2^+ve^ (75.31% cells per 4 sections), GPR88^+ve^ (69.35% cells per 4 sections) and Gabrg1^+ve^ (62.32% cells per 4 sections) cells with itch activated Fos^+ve^ cells in the CeA (Fig. 6d), which were shown to be significant enriched in itch-TRAPed tdTomato^+ve^ cells. To confirm whether this overlap is specific only to the enriched gene cluster, we also assessed the overlap between itch activated Fos^+ve^ cells in the CeA and cluster of genes with significantly lower expression in the itch-TRAPed neurons observed from RNAseq data. We find their partial overlap of Fos^+ve^ cells in the CeA (Fig. 6e) with cells that express OPRM1^+ve^ (26.47% cells per 4 sections) Penk^+ve^ (30.77% cells per 4 sections) and Chrm1^+ve^ (36.50% cells per 4 sections). These results confirm our findings of differentially expressed genes in itch-activated CeA cells and suggesting that further mining of these sequencing data by the community will reveal important new findings related to itch and its comorbidities.

**Fig 6.**
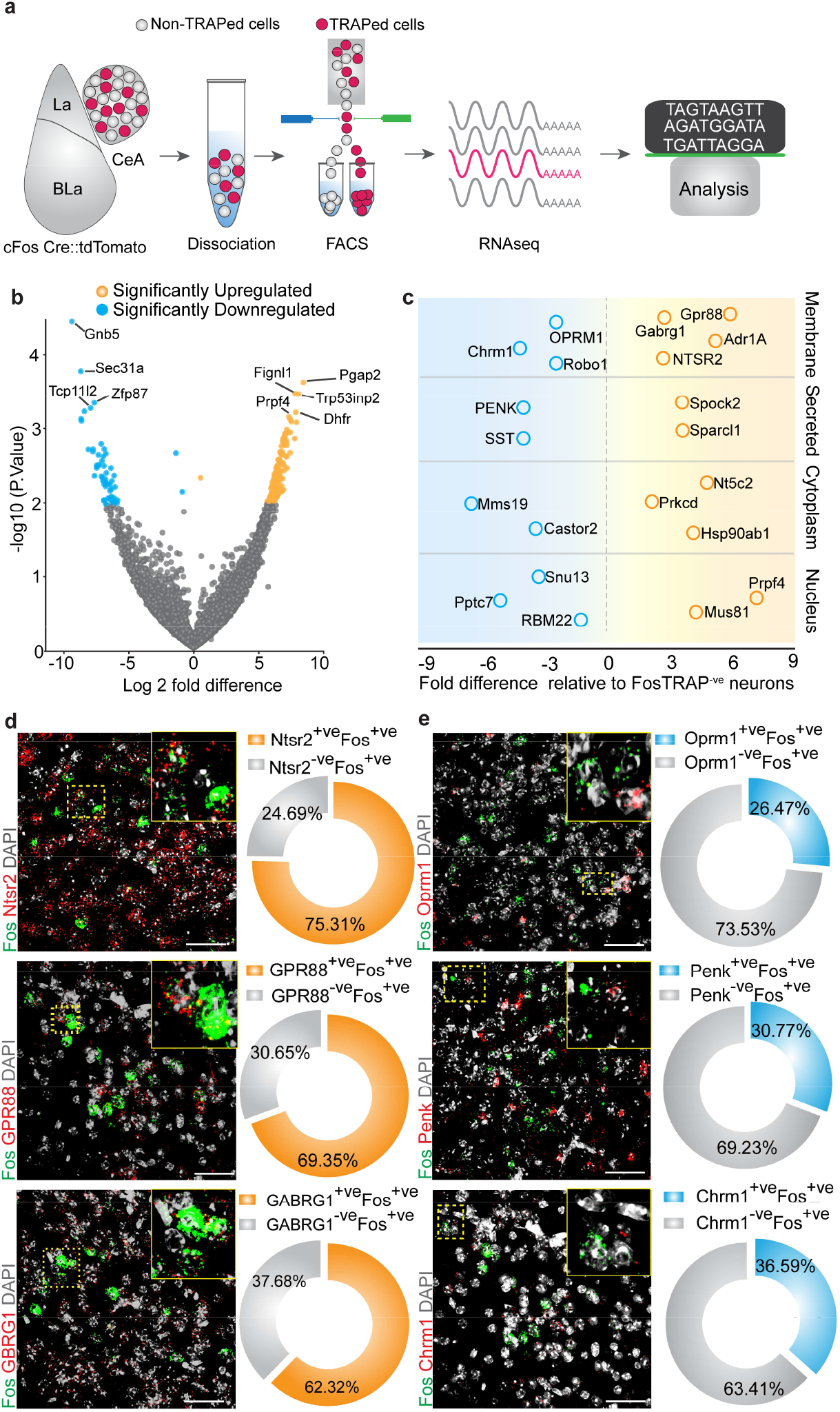
Cell-type specific transcriptomic profiling of itch-activated CeA cells. (a) Experimental workflow outlining fluorescence-activated-cell-sorting (FAGS) of the FosTRAPped tdTomato+ve and tdTomato-ve CeA neurons for whole-cell transcriptomics analyses. (b) Volcano plot of Log2-fold change (x axis) and p values (y axis) showing the transcripts that are differentially expressed in the itch-TRAPed tdTomato+ve CeA cells. Significantly differentially expressed genes are color coded and genes that have P <= 0.001 are indicated on the plot. (c) Candidate genes identified by fold change in expression of genes in significantly enriched KEGG pathways from the FosTRAPped tdTomato+ve CeA cells. (d) Multiplexed FISH was used in validating the expression of NTSR2, GPR88 and GABARG1 in itch activated Fos+ve CeA cells. We observed considerable overlap between NTSR2+ve (75.31% cells), GPR88+ve (69.35%) and Gabrg1+ve (62.32% cells) cells with itch activated Fos+ve cells in the CeA. (e) Multiplexed FISH was used to verify the overlap of OPRM1, Penk and Chrm1 in itch activated Fos+ve CeA cells. We find their partial overlap of Fos+ve cells in the CeA with cells that express OPRM1+ve (26.47% cells) Penk+ve (30.77% cells) and Chrm1+ve (36.50% cells). Right corner of each image shows magnification of the inset (yellow box).

## Discussion

While there is robust evidence demonstrating the role spinal cord circuits play in driving itch behaviors (Bautista et al., 2014; Han and Dong, 2014; LaMotte et al., 2014; Ross et al., 2010; Sun et al., 2009), the identity and properties of neural circuits in the brain that coordinate pruritic behaviors are still poorly understood. Neural circuits in the central amygdala are implicated in pruritic process (Chen et al., 2016; Mu et al., 2017), but cells and circuits that can alter pruritic processing in the CeA has been unclear. The CeA is well known to regulate a wide variety of aversive (Carrasquillo and Gereau, 2007; Ciocchi et al., 2010; Crock et al., 2012; Ehrlich et al., 2009; Haubensak et al., 2010; Tovote et al., 2016) and appetitive behaviors (Cai et al., 2014; Carter et al., 2013; Douglass et al., 2017; Hardaway et al., 2019; Kim et al., 2017; Robinson et al., 2014; Warlow et al., 2020). Here, we propose a cellular and circuit framework of the CeA in pruritis and its associated affect modulation. By gaining genetic access to neurons that are active specifically during itch, we were able to selectively identify a diverse repertoire of sensory and aversive behavioral responses mediated by itch-responsive CeA neurons. Activation of itch-responsive neurons in right or left CeA is sufficient to recapitulate spontaneous itch behaviors, potentiate chloroquine-evoked itch and produce aversive and anxiety-related behaviors. Furthermore, inhibiting these neurons is sufficient to attenuate ongoing itch and block its associated aversive component. Our findings reveal the presence of an itch-responsive neuronal population in the CeA that is necessary and sufficient to drive itch-related sensory and affective behaviors. We used both optogenetics and chemogenetics in a complementary manner to confirm these results. Conceptually, our findings seem to reconcile prior work (Chen et al., 2016; Mochizuki et al., 2020; Papoiu et al., 2014; Sun et al., 2009; Vierow et al., 2015) suggesting a pivotal role of CeA neurons in pruritic processing. The CeA is known to be involved in threat detection, and to produce adaptive responses when organisms encounter threatening conditions (Fadok et al., 2018; Grundemann and Luthi, 2015; LeDoux and Daw, 2018). These itch responsive CeA neurons could be a gateway in controlling itch and its associated affective component.

Activation of itch-TRAPed CeA neurons elicits aversive behaviors like place aversion and anxiety like behaviors, while inhibition of itch-TRAPed CeA neurons produces robust blockade of place aversion to chloroquine. It is well established that the CeA mediates stress induced anxiety behaviors (Botta et al., 2015; Kalin et al., 2004; Li et al., 2017; Weera et al., 2020), but our results show that silencing itch-TRAPed CeA neurons does not have any effect on basal anxiety like behavior. Future experiments should address whether these neurons are capable of attenuating anxiety during a variety of threats or stressors. Our findings also imply that itch-activated CeA neurons may also be involved to some extent in regulating nociceptive transmission. Itch-activated CeA neurons that drive pruritic processing could represent a subset or overlapping populations that also process nociceptive information. Our additional experiments also suggest that the CeA itch responsive neurons do not have any effect on feeding, reward seeking, motor and freezing behaviors. It is possible these behaviors are driven by distinct molecularly defined cell types in the CeA (Cai et al., 2014; Carter et al., 2013; Ciocchi et al., 2010; Douglass et al., 2017; Ehrlich et al., 2009; Hardaway et al., 2019; Haubensak et al., 2010; Kim et al., 2017; Robinson et al., 2014; Tovote et al., 2016; Warlow et al., 2020) that are distinct from the itch-activated CeA neurons we have studied here.

We found that itch-activated CeA neurons send functional outputs to the vPAG, and activating these CeA→vPAG projections is sufficient to drive scratching. These findings are consistent with recent reports (Gao et al., 2019; Samineni et al., 2019) showing that activating PAG^Vglut2^ and PAG^Tac1^ neurons produces robust scratching in mice. Based on the results from the present study and reconciling with prior work, a reasonable hypothesis is that information from itch-activated CeA neurons promotes scratching via disinhibition of the PAG^Tac1^ output neurons. As activating these PAG populations does not elicit freezing or escape behaviors, it is possible that PAG neurons that process pruritic information may be distinct from the ones that drive freezing, escape or nociceptive behaviors. Based on our findings, pruritic information arriving in PAG neurons originates at least in part via a CeA itch responsive neuronal population, and contributes to the processing and generation of adaptive responses to pruritis. Our RNAseq data show that CeA neurons are GABAergic, and anatomical tracing data indicate that CeA neurons that project to the PAG are GABAergic inhibitory populations (Supplementary Fig. 9d). Recent work shows that CeA GABAergic neurons that project to vlPAG can elicit freezing, escape (Haubensak et al., 2010; Tovote et al., 2016), hunting (Han et al., 2017), sleep (Snow et al., 2017) and nociception (Avegno et al., 2018; Li and Sheets, 2018; Yin et al., 2020). In our work, we found that the itch-activated CeA→PAG projections drive pruritic behaviors without evoking freezing or escape behaviors, suggesting that these FosTRAPed CeA→PAG projections are distinctly tuned to elicit scratching. The cellular and molecular identities that distinguish these projections from other CeA neurons is yet to be identified (Steinberg et al., 2020). A critical task in the future will be to identify and characterize how these different modalities of information are differentially processed via these projections and how the postsynaptic neurons in the PAG differentiate this information to transform it into behavioral output. Here we establish a critical role for inhibitory projections from CeA to PAG in pruritic regulation. Further studies will be required to more fully understand the mechanisms by which this projection drives itch and associated affective behaviors.

The CeA is highly molecularly heterogeneous region that is known to express a diverse array of neuropeptides, receptors and cellular machinery that are unique to CeA in integrating and orchestrating neuromodulatory functions (Kim et al., 2017; Zirlinger and Anderson, 2003; Zirlinger et al., 2001). To understand the unique genetic identity of CeA neurons that regulate pruritic behaviors, we performed RNAseq of itch-activated CeA neurons by TRAPing these neurons with pruritic stimuli. These activity-dependent RNA-Seq data show extensive molecular programs that are selectively enriched in the itch responsive neurons relative to other cells in CeA. We observe significant enrichment of NTSR2, GPR88 and Gabrg1 in itch-activated neurons, and relatively lower expression of OPRM1, PENK and Chrm1, suggesting that these genes could be critical candidates in regulating pruritis and its associated anxiety. In our analysis we have followed up on 6 candidate genes, but the dataset we have generated here will be an immensely valuable resource to the neuroscience community interested in the role of CeA neurons in modulation of sensory and affective behaviors.

## Methods

### Animals

All experiments were conducted in accordance with the National Institute of Health guidelines and with approval from the Animal Care and Use Committee of Washington University School of Medicine. Mice were housed on a 12-hour light-dark cycle (6:00am to 6:00pm) and were allowed free access to food and water. All animals were bred onto C57BL/6J background and no more than 5 animals were housed per cage. Male littermates between 8-12 weeks old were used for experiments. *FosCreERT2 mice (B6.129(Cg)-Fostm1.1(cre/ERT2)Luo/J; stock #21882, Ai9-tdTomato mice (B6.Cg-Gt(ROSA)26Sortm9(CAG-tdTomato)Hze/J; stock #007909,* Vgat-ires-Cre (*Slc32a1^tm2Lowl^*; *stock #028862.*), Vglut2-ires-Cre (*Slc17a6^tm2Lowl^*; stock # 028863.) and C57BL\6J mice were purchased from Jackson Laboratories and colonies were established in our facilities. For all the behavioral experiments heterozygous cFos-Cre male mice were used, for Cfos co-staining and sequencing experiments were performed on heterozygous cFos-Cre male mice crossed to homozygous Ai9mice from Jackson Laboratory. Litters and animals were randomized at the time of assigning experimental conditions for the whole study. Experimenters were blind to treatment and genotype.

### Viral constructs

Purified and concentrated adeno-associated viruses coding for Cre-dependent hM3Dq-mCherry(rAAV5/hSyn-DIO-hM3Dq-mCherry; 6 × 10^12^ particles/ml, Lot number: AV4495c and Lot date: 02/23/2012) and hM4D-mCherry (rAAV5/hSyn-DIO-hM4Di-mCherry; 6 × 10^12^ particles/ml, Lot number: AV4496c and Lot date: 11/20/2012), control mCherry (rAAV5/hSyn-DIO-mCherry; 3.4 × 10^12^ particles/ml, Lot number: AV5360 and Lot date: 04/09/2015), ChR2-eYFP (rAAV5-DIO-ChR2-eYFP; 4.8 × 10^12^ particles/ml, Lot number: AV4313Y and Lot date: 04/21/2017) and control eYFP (rAAV5-DIO-eYFP; 3.3 × 10^12^ particles/ml, Lot number: AV4310i and Lot date: 07/21/2016) was used to express in the FosCreERT2 mice. Helper virus, AAV1-EF1α-FLEX-TVAmCherry (rAAV5/ EF1α-FLEX-TVAmCherry; 4 × 10^12^ particles/ml) and AAV1-CAG-FLEX-RG (rAAV5/ CAG-FLEX-RG; 3 × 10^12^ particles/ml) were mixed at a ratio of 1:3 and then injected into the vPAG. Three weeks later, EnvA G-deleted Rabies-GFP (3.9 × 10^9^ particles/ml) was injected in the vPAG. All vectors except Rabies virus were packaged by the University of North Carolina Vector Core Facility. Rabies virus was purchased from Salk Gene Transfer, Targeting and Therapeutics Core. All vectors were aliquoted and stored in −80°C until use.

### Stereotaxic Surgeries

Mice were anesthetized with 1.5 to 2.0% isoflurane in an induction chamber using isoflurane/breathing air mix. Once deeply anesthetized, mice were secured in a stereotactic frame (David Kopf Instruments, Tujunga, CA) where surgical anesthesia was maintained using 2% isoflurane. Mice were kept on a heating pad for the duration of the procedure. Preoperative care included application of sterile eye ointment for lubrication, administration of 1mL of subcutaneous saline and surgery site sterilization with iodine solution. A small midline dorsal incision was performed to expose the skull and viral injections were performed using the following coordinates: CeA, −1.24 mm from bregma, +/− 2.8 mm lateral from midline and 4.5 mm ventral to skull. Viruses were delivered using a stereotaxic mounted syringe pump (Micro4 Microsyringe Pump Controller from World Precision Instruments) and a 2.0uL Hamilton syringe. Injections of 75-100 nL of the desired viral vectors into the area of interest were performed at a rate of 1uL per 10 minutes. We allowed for a 10-minute period post injection for bolus diffusion before removing the injection needle. Postoperative care included closure of the cranial incision with sutures and veterinary tissue adhesive, and application of topical triple antibiotic ointment to the incision site. Animals were monitored while on a heating pad until they full recovery from the anesthetic.

### Cannula Implantation

The surgical protocol was the same as described above for viral injections. Fiber optic implants were fabricated using zirconia ferrules (Thorlabs) and from 100μm diameter fibre (0.22 numerical aperture (NA), Thorlabs). Fiber optic cannulas (length 5mm) were implanted at the CeA and the PAG and fixed to the skull using two bone screws (CMA anchor screws, #7431021) and dental cement. The following coordinates were used for implantation: CeA, −1.24 mm from bregma, +/− 2.8 mm lateral from midline and 4.25 mm ventral to skull and the PAG, −4.84 mm from bregma, +/− 0.5 mm lateral from midline and 2.7 mm ventral to skull. Mice were allowed to recover for 14 days before behavioral analysis. Animals in which cannulas placement missed the CeA or vlPAG target were excluded from the study.

#### Chemogenetic manipulation

For chemogenetic control of CeA FosTRAPped neurons, cFos-Cre mice were injected with Cre dependent control mCh, hM3Di or hM4Dq viruses. DREADD constructs used in this study were validated previously in our lab for their functional expression in the PAG, including their ability to increase (hM3Dq) or decrease (hM4Di) neuronal firing in slices from animals expressing these viral constructs (Samineni et al., 2017a). Three weeks later mice were injected with 4-OHT to express Cre dependent DREADDs, clozapine N-oxide (CNO, BML-NS105 from Enzo life sciences) was injected 30 min before doing behavioral experiments and data were collected between 30 min- and 2-hours post-injection. All baselines for pruritic responses were recorded 3 weeks after the FosTRAP and one week prior to the CNO administration. We used 5 mg/kg CNO as a dose of CNO and showed no signs of behavioral changes in control vector-expressing animals.

#### Optogenetic manipulations

For all the behavioral experiments mice were acclimated to tethered fibers for 5 days before initiation of the experiments. Mice were habituated to tethering with lightweight patch cables (components: Doric lenses) that are connected to a laser (Shanghai laser, 475nm). To prevent impediment of movement from the tethered cables, we coupled patch cables to an optical commutator (Doric Lenses). An arduino was programed and connected to the laser to deliver 5, 10, 20 and 30 Hz (5-ms width, 10mW/mm^2^) photostimulation in FosCre mice.

#### Activity-dependent FosTRAP labeling

##### 4-hydroxytamoxifen preparation and delivery

We dissolved 10 mg of 4-OHT (Sigma, Cat# H6278-10MG) in 500ul ethanol (100%) (20mg/ml stock) first by vortexing and then sonicating.,We then add autoclaved corn oil (1:4) to dissolve 4-OHT (previously heated to 45^0^C) to 5mg/mL and sonicate until solution cloudiness clears. As a final step vacuum centrifuge for 10 min to evaporate the alcohol from the final injection solution. Male FosTRAP (FosCreER+/−, FosCreER+/−, Ai9+/−) mice were used. Mice were single housed and gently handled for 7-10 days prior to the experiment to minimize the unwanted labelling of neurons associated with stress of handling. On the experiment day mice were given 4OHT 20mg/kg in their homecage environment. 60 Min post 4-OHT, we injected either saline or chloroquine (200ug/50ul) subcutaneously in the nape of the neck to TRAP neurons that are activated by pruritic stimuli. In FosCreER+/−, Ai9+/− mice, robust tdTomato expression was seen 1 week post TRAPing. In the FosCreER+/− mice injected with the optogenetic or chemogenetic constructs, robust labelling was seen 4 weeks post TRAPing.

#### Pruritic agent induced scratching behaviors

As previously described by our group (O’Brien et al., 2013; Valtcheva et al., 2015), the nape of the neck of mice was shaved one day prior to experiments. Mice were then placed in clear plexiglass behavioral boxes for at least two hours for acclimation. For chemogenetic manipulations, CNO was administered before placing the mice in the plexiglass behavioral boxes and chloroquine (200μg/50μl, nape of the neck) induced scratching behavior was performed 90 min after the CNO administration.

#### Pain behavior assessment

Mechanical sensitivity was measured by counting the number of withdrawal responses to 10 applications of von Frey filaments (North Coast Medical, Inc, Gilroy, CA; 0.02, 0.08, 0.32 and 1.28 g von Frey filaments) to both hindpaws as described (Samineni et al., 2017c). Each mouse was allowed at least 15 seconds between each application and at least 5 minutes between each size filament. Animals were acclimated to individual boxes on a plastic screen mesh for at least one hour before testing. The Hargreaves test was performed to evaluate heat sensitivity thresholds as previously described (Samineni et al., 2017a). Briefly, we measured latency of withdrawal to a radiant heat source (IITC Life Science, Model 390). We applied the radiant heat source to both hindpaws and measured the latency to evoke a withdrawal. Three replicates were acquired per hindpaw per mouse and values for both paws were averaged.

#### Open-Field test (OFT)

Before testing, mice were habituated to the test room in their home cages for 2 h. Control and mice injected with either hM3Dq, hM4Di or ChR2 in the CeA were then placed in the open field during individual trials and allowed to freely explore after the experimenter exited the room (behaviors were video recorded). Open field locomotor activity was assessed in a square enclosure (55 × 55 cm) within a sound attenuated room for 30 min (Shin et al., 2017). Total distance traveled and movements were video recorded and analyzed using Ethovision XT (Noldus Information Technologies, Leesburg, VA).

#### Elevated Zero maze (EZM)

Anxiety was measured in low light conditions (~20 lux) using a modified zero maze (Stoelting Co., Wood Dale, IL) placed 70 cm off of the ground and consisting of two closed sections (wall height, 30 cm) and two open sections (wall height, 1.3 cm) on a circular track (diameter of track, 60 cm)(Montana et al., 2011). On the experiment day mice were habituated to testing room for 1 h before beginning of the behavioral session. For hM3Dq and hM4Di injected mice 60 minutes after CNO injection, mice were placed individually at the intersection of the closed/open area of the zero maze for a 600-s trial. For Chr2 and eYFP FosTRAP mice, mice were connected to the fiber optic and placed at the intersection of the closed/open area of the zero maze for a 600-s trial. Mice received 20 Hz (5ms width) photostimulation for the duration of the EZM trial. Movement during the trial was video recorded using digital camera (Floureon HD) mounted on the ceiling of the room. Total distance traveled, number of entries into open sections, and time spent in the open sections was scored were video recorded and analyzed using Ethovision XT (Noldus Information Technologies, Leesburg, VA).

#### Realtime place aversion testing (RPA)

Place aversion was tested in a custom designed two compartment chamber (52.5 × 25.5 × 25.5 cm) with a layer of corn cob bedding (Shin et al., 2017). Each mouse was placed in the neutral area of the chamber and given free access to roam across both chambers. Activity was continuously recorded through a video camera for a period of 20 min. Entry into light-paired chamber triggered constant photostimulation at either 5Hz, 10Hz, 20Hz or 30Hz (473 nm, 5 ms pulse width, ~10 mW light power). Entry into the other chamber terminated the photostimulation. Photostimulation was counterbalanced across mice. “Time-in-chamber” and heat maps were generated for data analysis using Ethovision XT software (Noldus, Leesburg, VA).

#### Conditioned place aversion (CPA)

CPA was performed using an unbiased, counterbalanced three-compartment conditioning apparatus as described (Land et al., 2009). Each chamber had a unique combination of visual, properties (one side had black and white vertical walls, whereas the other side had black and white horizontally striped walls). On the pre-conditioning day (day 1), mice were allowed free access to all three chambers for 20 min. Behavioral activity in each compartment was monitored and recorded with a video camera and analyzed using Ethovision 8.5 (Noldus) or ANY-Maze software. Mice were randomly assigned to saline and chloroquine compartments and received a saline injection (50μl) in the nape of the neck and on the mouse caudal back, in the morning and a chloroquine injection (200μg/50μl) in the nape of the neck and on the mouse caudal back in the afternoon, at least 4 h after the morning training on 3 consecutive days (Day 2, 3 & 4). To enhance the association of chloroquine induced scratching behavior with the paired chamber, we administered chloroquine and left the mice in their holding cage for 4 min, then placed them in the paired chamber during the time of the peak scratching response (20 min in the chamber). To assess for place aversion, the mice were then allowed free access to all three compartments on day 5 for 30 minutes (Tzschentke, 2007). Scores were calculated by subtracting the time spent in the chloroquine-paired compartment, post-test minus the pre-test. To test the effect of DREADD hM4Di activation on chloroquine-induced place aversion, mice injected with AAV5-DIO-hM4Di–mCherry and AAV5-DIO-mCherry were allowed free access to all three chambers for 30 min on the pre-conditioning day (day 1). On day 2, 3 & 4, both cohorts received a saline injection (50μl) in the nape of the neck and on the mouse caudal back, and this chamber was paired with systemic saline injection 1 h before they were placed in the compartment in the morning and a chloroquine injection (200μg/50μl) in the nape of the neck and on the caudal back and this chamber was paired with systemic CNO injection 1 h before they were placed in the compartment in the afternoon. To test the effect of DREADD hM4Di activation on chloroquine-induced place aversion, the mice were allowed free access to the three compartments on day 5 for 30 minutes. Scores were calculated by subtracting the time spent in the chloroquine-paired compartment, post-test minus the pre-test.

#### Operant conditioning

Mice are food deprived to reach 90% of their body weight and trained to nose poke for sucrose pellets for 7 days during daily 60 min sessions in a modular test chamber (Med Associates) on a fixed-ratio 1 (FR1) schedule of reinforcement as previously described by (Seo et al., 2016; Shin et al., 2017). A correct nose poke response in the active hole resulted in a sucrose pellet delivery where an incorrect nose poke within the inactive hole resulted in no sucrose pellet. On the experiment day, mice were administered CNO followed by a 60 min operant self-stimulation session. To determine if DREADD manipulation of FosTRAPed CeA neurons has any effect on the fixed-ratio 1 (FR1) schedule of reinforcement, mice were given free access to nose poke the ports, three successive nose pokes (FR3) to the active portal rewarded the mouse a sucrose pellet delivery where an incorrect nose poke within the inactive hole resulted in no sucrose pellet. On the experiment day, mice were administered CNO followed by a 60 min operant self-stimulation session, to determine if DREADD manipulation of FosTRAPed CeA neurons has any effect on fixed-ratio 3 (FR3) schedule of reinforcement.

#### Feeding behavior

Mice were given free access to a novel empty cage prior to the experiment day. Mice were food-deprived overnight prior to the experiment day (Cai et al., 2014). Mice were reintroduced into the same empty cage they had access to the prior day but with food pellets and allowed to feed freely for 20 min on the experiment day. At the end of the session, weight of the food pellet and the food debris left on the cage floor was measured to calculate the food intake. To determine whether FosTRAPed CeA neurons modulate feeding behaviors, mice were injected with CNO 60 min before the feeding test. Feeding tests were performed between 2 p.m. and 7 p.m.

#### Fiber Photometry

For *in vivo* calcium imaging of CeA GABAergic neurons, we injected the CeA of Vgat-Cre mice with Cre-dependent GCaMP6s (AAV-DJ EF1a-DIO-GCaMP6s, 3 × 10^13^ particles/ml, Stanford vector core). Fiber optic probes were unilaterally implanted above the right CeA (−1.24 mm from bregma, +/− 2.84 mm lateral from midline and 4.4 mm ventral to skull). After 4 weeks of viral expression, mice were handled and acclimated by tethering as will occur during imaging sessions, for 7 days in the test behavioral chamber. On the test day, mice were habituated with the tethered fiber optic patch cord (0.48NA, BFH48-400, Doric lenses) in the test chamber (15 × 15cm) for 60 min and then injected with chloroquine (200ug/50ul) in the nape of the neck and recordings were performed.

A fiber optic patch cord was used to connect to the fiber implant and deliver light to excite and record the GCAMP signal using a custom-built fiber photometry rig, built with some modifications to previously described specifications (Cui et al., 2013). Fluorescence excitation was provided by two LEDs at 211 and 537 Hz to avoid picking up room lighting (M405FP1, M470F1; LED driver: LEDD1B; Thorlabs). Light was bandpass filtered (FMC1+(405/10) −(475/28)_(525/45)_FC), Doric lenses) and delivered to the CeA to excite GCaMP6s. The emitted light was bandpass filtered (FMC1+_(405/10)-(475/28)_(525/45)_FC), Doric lenses) and sent to a photoreceiver to detect the signal (Newport, 2151). The signal from the photoreceiver was recorded using a RZ5P real-time processor (TDT). Data were acquired at 10 kHz and then demodulated at 211 and 537 Hz. The demodulated signal was then low-pass filtered (4 Hz) in a custom MATLAB script. The extracted 405 nm signal was then scaled to fit the GCaMP signal for the recording session. To isolate the movement-corrected GCaMP signal from channel, we subtracted the signal at 405 nm from the 475 nm GCaMP signal. dF/F was obtained by dividing the final signal with its mean value. Behavioral event timestamps associated with chloroquine evoked scratching behavior were scored and aligned with GCaMP signal in the MATLAB script to create pre- and peri-stimulus time bins. To obtain pre- and peristimulus chloroquine evoked scratching events, if the scratching events happened close to each other (in a 30 second window) they we combined and scored as one bout. Z-score was obtained by subtracting the mean of the GCaMP signal from the bin value of the GCaMP signal and dividing it with the standard deviation of the bin value of the GCaMP signal.

#### Acute slice electrophysiology

To determine the functional effects of chemogenetic manipulations in the itch FosTRAPped CeA neurons, we performed targeted whole-cell patch-clamp recordings in acute coronal slices from cFos-Cre mice expressing either hM3Dq or hM4Di receptors as previously described(Samineni et al., 2017a). Mice used for electrophysiology and behavioral studies were between 8-16 weeks of age. Three weeks after viral injections, we performed itchTRAP and waited 3 weeks for expression of hM3Dq or hM4Di in the CeA. Coronal slices containing the CeA were prepared and CeA neurons were visualized through a 40x objective using IR-DIC microscopy on an Olympus BX51 microscope, and mCherry+ cells were identified using epifluorescent illumination with a green LED (530 nm; Thorlabs), coupled to the back-fluorescent port of the microscope. Whole cell recordings of itch FosTRAPed CeA neurons expressing hM3Dq-mCherry and hM4Di-mCherry were performed using a Heka EPC 10 amplifier (Heka) with Patchmaster software (Heka). Following stable 5 min whole-cell recordings (baseline), the effects of either hM3Dq or hM4Di receptor activation on cellular excitability was isolated by blocking AMPA/KARs (10 μM NBQX, Abcam), NMDARs (50 μM D-APV, Abcam), GABAARs (100 μM picrotoxin, Abcam), and GABABRs (50 μM baclofen, Abcam), and aCSF solution containing 10 μM CNO added to the antagonist cocktail above was bath applied to the brain slice.

#### Immunohistochemistry

Adult mice were deeply anesthetized using a ketamine/xylazine cocktail and then perfused with 20ml of phosphate-buffered saline (PBS) and 4% paraformaldehyde (weight/volume) in PBS (PFA; 4 °C). *For Fos staining:* To determine the causal contribution of the CeA neuron in itch processing, we gave chloroquine to the nape of the neck and 90 min later mice were perfused. To verify whether itch TRAPped tdTomato+ CeA neurons are faithfully TRAPed to pruritic stimulus and rule out non-specific labeling, one week after the TRAP, we gave a second chloroquine injection and 90 min later mice were perfused. Brains were carefully removed, post fixed in 4% PFA overnight and later cryoprotected by immersion in 30% sucrose for at least 48 hrs. Tissues were mounted in OCT while allowing solidification of the mounting medium at −80 °C. Using a cryostat, 30μm tissue sections were collected and stored in PBS1x 0.4% sodium azide at 4 °C. After washing the sections in PBS1x, we blocked using 5% normal goat serum and 0.2% Triton-X PBS 1x for one hour at room temperature. Primary antibodies against mCherry (Mouse, Clontech, 632543; 1/500), GFP (rabbit polyclonal anti-GFP, Aves A11122; 1/500) and cFos, (rabbit monoclonal anti-phospho-cFos, Cell Signaling Ser32 D82C12; 1:2000) were diluted in blocking solution and incubated overnight at 4 °C. After three 10-minute washes in PBS1x, tissues were incubated for one hour at room temperature with secondary antibodies (Life Technologies: Alexa Fluor488 donkey anti rabbit IgG (1/500); Alexa Fluor 488 goat anti rabbit (1/500); Alexa Fluor 555 goat anti mouse (1/500) Alexa Fluor 555 goat anti rabbit (1/500)) and Neurotrace (435/455nm, 1/500) at room temperature. Three PBS1x washes followed before sections were mounted with Vectashield (H-1400) hard mounting media and imaged after slides cured. Images were obtained on a Nikon Eclipse 80i epifluorescence microscope.

#### Tissue preparation for RNAseq and Fac Sorting

Animals (8-10-week-old, 7-10 days post TRAP) were used for this experiment to ensure robust Ai9 reporter expression, while assuring fully developed brains. RNA-seq of the TRAPped neurons was performed using protocols modified from prior published work to improve neuronal survival (Arttamangkul et al., 2006; Guez-Barber et al., 2012; Hempel et al., 2007). Animals were anesthetized with ketamine cocktail, perfused with aCSF (124 mM NaCl, 24 mM NaHCO_3_, 12.5 mM glucose, 2.5 mM KCl, 1.25 mM NaH_2_PO_4_, 2 mM CaCl_2_, 1 mM MgCl_2_, 5mM HEPES, pH 7.4, 300-310 mOsm) and decapitated for brain removal. The brains were allowed to rest in cold oxygenated (95% O2/5% CO_2_) aCSF and then sliced coronally using a vibratome (Leica VT1000 S). Brain slices (400-μm thick sections) were collected and kept in cold oxygenated aCSF. Tissues were micro-dissected under a microscope (Leica S9i) using a reusable 0.5mm biopsy punch (WPI 504528). HBSS+H and Papain solution (45U, Worthington, Lakewood, NJ)) was incubated for 5min at 37 degrees, followed by the addition of tissue punches for 10-15min. Tissue punches were then transferred to ice, and mechanical trituration of tissue punches was performed using ~600, 300 and 150 μm fire-polished Pasteur pipettes. The resulting cell suspension was then centrifuged at 5k RPM for 5min to obtain a pellet, and cells were re-suspended in fresh aCSF. This process happened twice to wash any remnants of Papain. Cells were ultimately resuspended for FACS sorting into cold oxygenated aCSF and kept on ice for the duration of the experiment.

Cell suspensions were kept cold throughout the FACS, and cells were sorted in aCSF. In order to determine gating criteria for selecting cell bodies while excluding debris, we performed FACS on fixed/permeabilized neurons stained with Neurotrace 435/455nm (Nissl stain). Samples were treated with 2% PFA for 20min, pelleted down for 5min at 5k RPM, and then resuspended in PBS1x 0.3% Triton X-100. This processed was done an additional time to get rid of any remnant of PFA. Cells were then resuspended in aCSF and incubated with Neurotrace 435/455 (ThermoFisher, #N211479) for FACS sorting. We gated for events which had high levels of Neurotrace, and then mapped these events in the scatter plot (forward scatter FSC vs. side scatter SSC). We were able to map events which had high Neurotrace expression to a small subset of events, which represent the population of cell bodies and not debris. In addition, this population was sensitive to PFA fixation and labeling with the nuclear staining DAPI or 7-AAD, which is characteristic of post-fixative dead cells. As for DAPI/7-AAD (dead) control samples, these were incubated in 2% PFA for 20min, pelleted down for 5min at 5k RPM, and then resuspended in PBS1x 0.3% Triton X-100, this processed was done an additional time to get rid of any remnant PFA. Cells were then resuspended in aCSF and incubated with DAPI (1:1000 dilution of 1mg/mL DAPI, Thermo Fisher, #62248 or 7-AAD 7-Aminoactinomycin D, A1310, Thermo Fisher) for FACS sorting. We performed control experiments to set the appropriate gates for florescence, Ai9 (tdtomato) expression. Negative control samples were obtained from c57BL6/J animals, while positive controls were obtained from Vgat Ai9. FosCre × Ai9 brains were used for isolation of the neuronal population of interest. The CeA was dissociated as previously described (Guez-Barber et al., 2012), and cells were sorted into a 96-well plate. Up to a maximum of 50 cells were sorted into one well filled with 9uL of Clontech lysis buffer (Single-cell lysis buffer 10x, #635013 Takara Bio) + 5% RNAse inhibitor (40U/ul, Promega RNAsin inhibitor N2511). Samples were then transferred to a tube for processing by our Genome Technology Access Center (GTAC) core facility. ds-cDNA was prepared using the SMARTer Ultra Low RNA kit for Illumina Sequencing (Takara-Clontech) per manufacturer’s protocol using the lysis buffer as substrate for the reaction. cDNA was fragmented using a Covaris E220 sonicator using peak incident power 18, duty factor 20%, cycles/burst 50, time 120 seconds. cDNA was blunt ended, had an A base added to the 3’ ends, and then had Illumina sequencing adapters ligated to the ends. Ligated fragments were then amplified for 15 cycles using primers incorporating unique index tags. Fragments were sequenced on an Illumina HiSeq-3000 using single reads extending 50 bases.

#### RNAseq

RNA-seq reads were aligned to the Ensembl top-level assembly with STAR version 2.0.4b. Gene counts were derived from the number of uniquely aligned unambiguous reads by Subread:featureCount version 1.4.5. Transcript counts were produced by Sailfish version 0.6.3. Sequencing performance was assessed for total number of aligned reads, total number of uniquely aligned reads, genes and transcripts detected, ribosomal fraction known junction saturation and read distribution over known gene models with RSeQC version 2.3.

All gene-level and transcript counts were then imported into the R/Bioconductor package EdgeR and TMM normalization size factors were calculated to adjust for samples for differences in library size. Ribosomal features as well as any feature not expressed in at least the smallest condition size minus one sample were excluded from further analysis and TMM size factors were recalculated to created effective TMM size factors. The TMM size factors and the matrix of counts were then imported into R/Bioconductor package Limma and weighted likelihoods based on the observed mean-variance relationship of every gene/transcript and sample were then calculated for all samples with the voom with quality weights function. Generalized linear models were then created to test for gene/transcript level differential expression. Differentially expressed genes and transcripts were then filtered for p-values less than or equal to 0.001.

The biological interpretation of the genes found in the Limma results were then queried for global transcriptomic changes in known Gene Ontology (GO) and KEGG terms with the R/Bioconductor packages GAGE and Pathview. Briefly, GAGE measures for perturbations in GO or KEGG terms based on changes in the observed log 2-fold-changes for the genes within that term versus the background log 2-fold-changes observed across features not contained in the respective term as reported by Limma. For GO terms with an adjusted statistical significance of FDR <= 0.05, heatmaps were automatically generated for each respective term to show how genes co-vary or co-express across the term in relation to a given biological process or molecular function. In the case of KEGG curated signaling and metabolism pathways, Pathview was used to generate annotated pathway maps of any perturbed pathway with an unadjusted statistical significance of p-value <= 0.05.

To find the most critical genes, the raw counts were variance stabilized with the R/Bioconductor package DESeq2 and was then analyzed via weighted gene correlation network analysis with the R/Bioconductor package WGCNA. Briefly, all genes were correlated across each other by Pearson correlations and clustered by expression similarity into unsigned modules using a power threshold empirically determined from the data. An eigengene was then created for each de novo cluster and its expression profile was then correlated across all coefficients of the model matrix. Because these clusters of genes were created by expression profile rather than known functional similarity, the clustered modules were given the names of random colors where grey is the only module that has any pre-existing definition of containing genes that do not cluster well with others. The information for all clustered genes for each module were then combined with their respective statistical significance results from Limma to determine whether or not those features were also found to be significantly differentially expressed. Raw and analyzed data can be found at GEO: GSE130268.

#### Fluorescence in situ hybridization (FISH)

C57BL/6 J mice were injected with chloroquine on the nape of the neck. Thirty minutes post chloroquine administration, mice were rapidly decapitated, brains were dissected and flash frozen in −50 °C 2-methylbutane and stored at −80 °C for further processing(Samineni et al., 2017b). Coronal sections of the brain corresponding to the CeA, were cut at 15 μM at −20°C and thaw-mounted onto Super Frost Plus slides (Fisher). Slides were stored at −80 °C until further processing. FISH was performed according to the RNAScope® 2.0 Fluorescent Multiple Kit v2 User Manual for Fresh Frozen Tissue (Advanced Cell Diagnostics, Inc.). Slides containing CeA sections were fixed in 4% paraformaldehyde, dehydrated, and pretreated with protease IV solution for 30 min. Sections were then incubated with target probes for mouse cFos (mm-Fos, catalog number 316921, Advanced Cell Diagnostics), Ntsr2 (mm-Ntsr2, catalog number 452311, Advanced Cell Diagnostics), GPR88 (mm-GPR88, catalog number 317451, Advanced Cell Diagnostics), Penk (mm-Penk, catalog number 318761, Advanced Cell Diagnostics), Gabarg1 (mm-Gabarg1, catalog number 501401, Advanced Cell Diagnostics), Oprm1 (mm-Oprm1, catalog number 315841, Advanced Cell Diagnostics), Chrm1 (mm-Chrm1, catalog number 495291, Advanced Cell Diagnostics). Following probe hybridization, sections underwent a series of probe signal amplification steps (AMP1–4) followed by incubation of fluorescent probes (Opal 470, Opal 570, Opal 670), designed to target the specified channel associated with the probes. Slides were counterstained with DAPI and coverslips were mounted with Vectashield Hard Set mounting medium (Vector Laboratories). Images were obtained on a Leica TCS SPE confocal microscope (Leica), and Application Suite Advanced Fluorescence (LAS AF) software was used for analyses. To quantify number of cFos+ve cells, we counted DAPI-stained nuclei that coexpress minimum of five cFos puncta as an cFos+ve cell. We did not include any cFos puncta that does not overlay on top of the DAPI-stained nuclei as part of our analysis.

#### Statistics

Throughout the study, researchers were blinded to all experimental conditions. Exclusion criteria for our study consisted on a failure to localize expression in our experimental models or off-site administration of virus or drug. At least 3 replicates measurements were performed and averaged in all behavioral assays. The number of animals used is indicated by the “N” in each experiment. When paired t Test was used for comparing paired observations, we evaluated for normality using the D’Agostino and Pearson omnibus normality test for all datasets. Therefore, only when normality could be assumed, we used a parametric test to analyze out data. If normality could not be assumed, a nonparametric test or a Wilcoxon matched pairs text was used to evaluate differences between the means of our experimental groups. Two-way ANOVA was used for comparing between different control and treatment groups. Bonferroni’s *post hoc* tests were used (when significant main effects were found) to compare effects of variables. A value of p <0.05 was considered statistically significant for all statistical comparisons.

## Acknowledgments

This work was funded by NINDS R01NS106953 and R01DK116178 to RWG, the Urology Care Foundation Research Scholars Program and Kailash Kedia Research Scholar Award and NIDDK Career development award (K01 DK115634) to VKS, NRSA F32 DK115122 to A.D.M. and the Medical Scientist Training Program (MSTP) Grant T32GM07200 and NINDS NRSA 5F31NS103472-02 to JGGR. We Kenneth M. Murphy and his lab for assistance with FACS. We thank Sherri Vogt for her assistance with mouse colony maintenance and genotyping. We would like to thank Dr. Jordan G McCall for helpful discussion with the manuscript and experimental help with slice electrophysiology; We also thank Daniel Castro and Adrian Gomez for their help with reward seeking experiments; We would like to thank all the Gereau lab members for their help with manuscript preparation.

## Author Contributions

V.K.S and R.W.G. designed the experiments; V.K.S., J.G.G., E.S.A, J.N.S, S.B.S. performed anatomical analyses; V.K.S. and J.G.G. performed behavior; C.P wrote the code for photometry analysis; G.E.G performed FACS for RNA-seq; E.T. Performed data curation, analysis and methodology for RNA-seq; B.A.C. performed slice electrophysiology; M.R.B. contributed reagents/resources; V.K.S., J.G.G., G.E.G., B.A.C., A.D.M, E.T., C.P, and R.W.G. analyzed the data; V.K.S., and R.W.G. wrote the manuscript with comments from all the authors.

## Code availability

Matlab code used in this study is available from the corresponding author upon request.

