## Supplementary figures for "Cellular, Circuit and Transcriptional Framework for Modulation of Itch in the Central Amygdala"

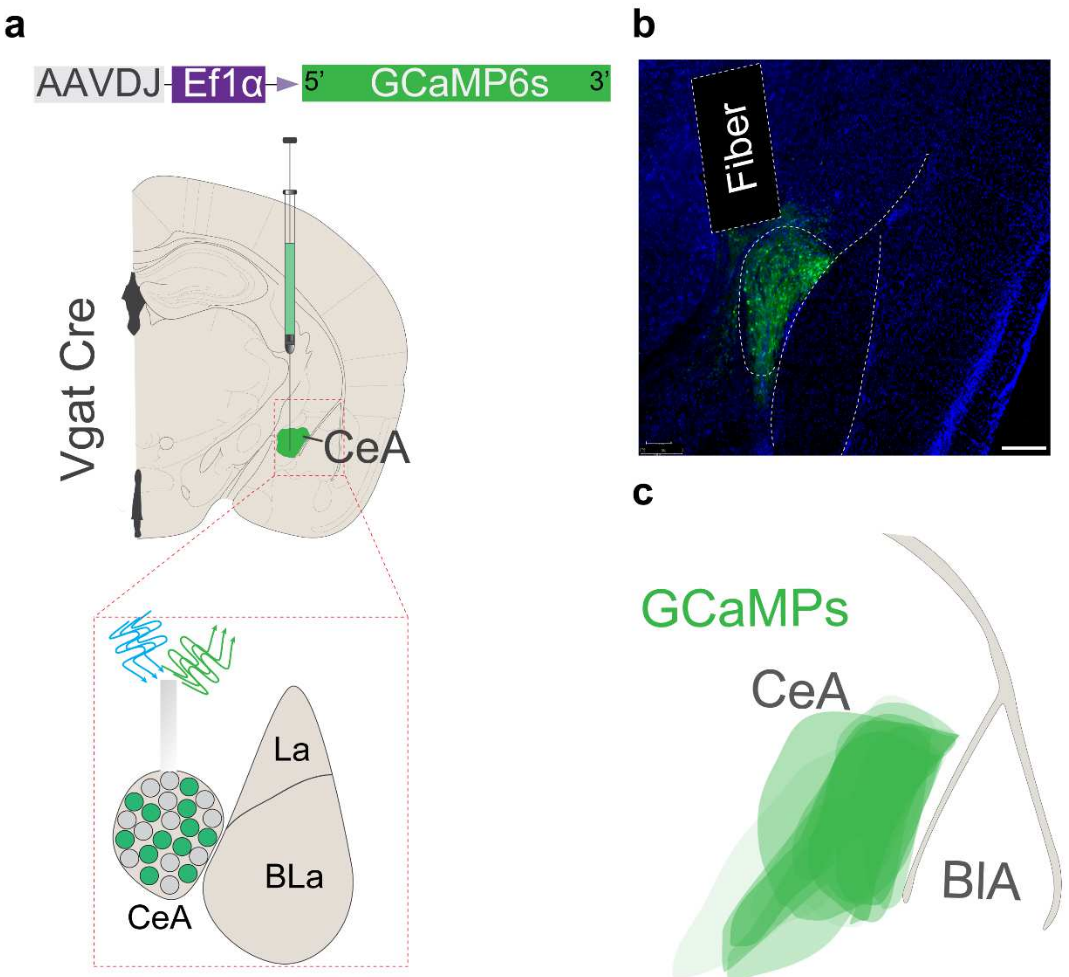


**Supplementary Fig 1. Anatomical location of the GCaMP6s-exprssing CeA neurons and fiber placements for imaging activity during itch behaviors.** (a) Scheme demonstrating viral injection strategy and fiber placement to record CeAVgat neural activity in response to chloroquine. (b) Representative image of the CeA of Vgat Cre mice in which AAV-DJ-DIO-GCaMP6s is expressed and fiber placement for photometry. Scale bar, 200 μm. (c) Illustration showing the CeA viral spread of AAV-DJ-DIO-GCaMP6S injection.

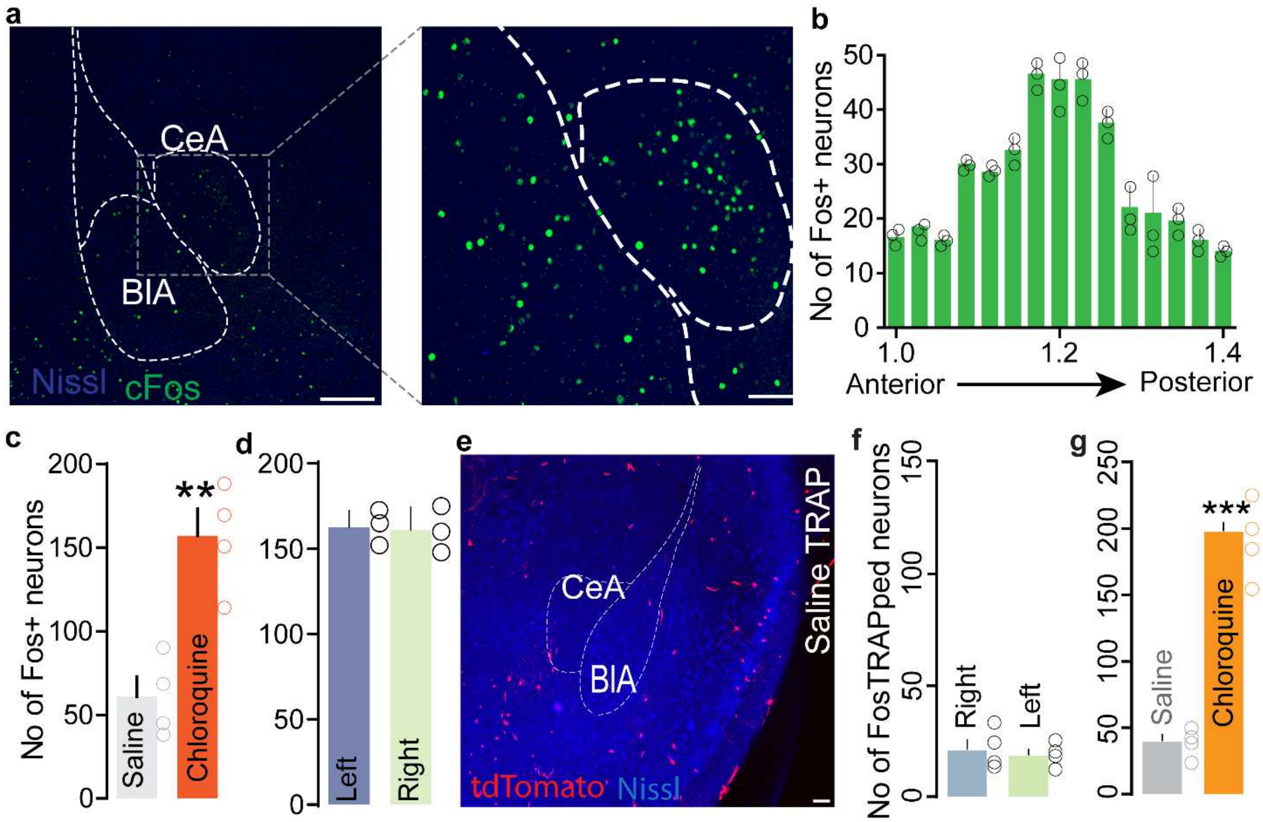


**Supplementary Fig 2. Anatomical location of cFos-expressing (itch-activated) neurons in the CeA following chloroquine injection in the nape of the neck.** (a) Representative sections showing cFos labeling at low and high magnification Scale bar 250 µm in left panel, 50 µm in the right panel. (b) Rostro-caudal distribution of CeA cFos+ve neurons in CeA. (c) Number of c-Fos+ve neurons in the CeA after administering either saline or chloroquine. n = 4 per group, t=4.801 df=6 ** $p < 0.01$. (d) Quantification of total number of c-Fos+ve neurons in the left and the right CeA after administering chloroquine stimuli. n = 3 per group, t test, t=0.01779, df=4, p = 0.9867. (e) FosTRAPing with saline produced very few tdTomato+ve cells. Scale bar, 75 µm. (f) Quantification of total number of saline FosTRAPped neurons in left and right CeA. n = 3 per group. t test, t=0.188, df=2, p =0.85. Scale bar, 75 µm. (g) FosTRAPing with chloroquine significantly increased the number of tdTomato+ve neurons compared to saline. n = 3 per group, t=3.155 df=6 *** $p < 0.001$.

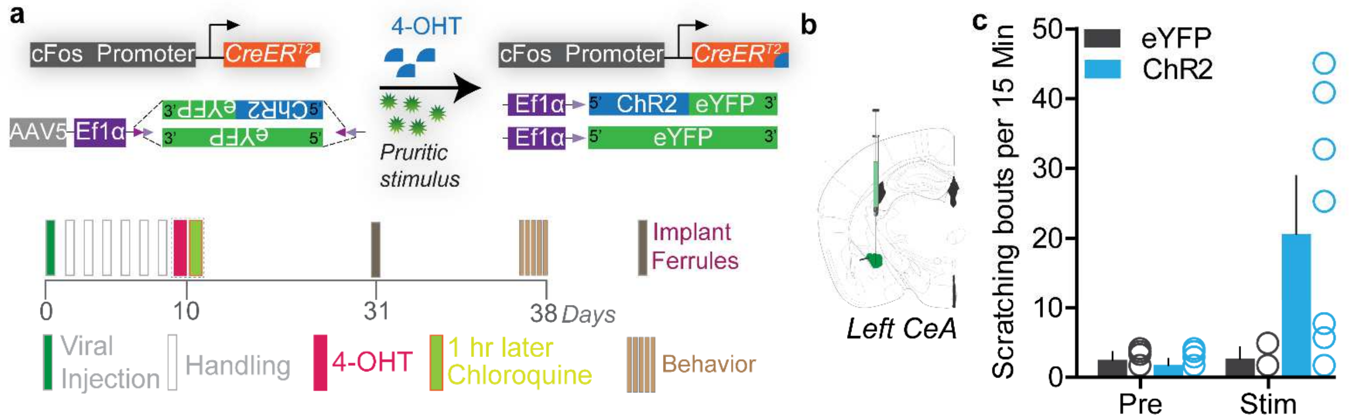


**Supplementary Figure 3. Optogenetic re-activation of itch-TRAPped neurons in the left CeA neurons promotes scratching.** (a) Viral strategy to selectively express ChR2 or eYFP in itch-activated neurons of the left CeA. Experimental timeline to FosTRAP ChR2/eYFP in CeA. (b) Viral strategy to selectively express optogenetic constructs in CeA FosTRAPed neurons. (c) Photostimulation of itch-activated neurons in the left CeA produces significant scratching in ChR2 FosTRAPed mice, but not in eYFP FosTRAPped controls. n = 6–8 per group, $p < 0.05$.

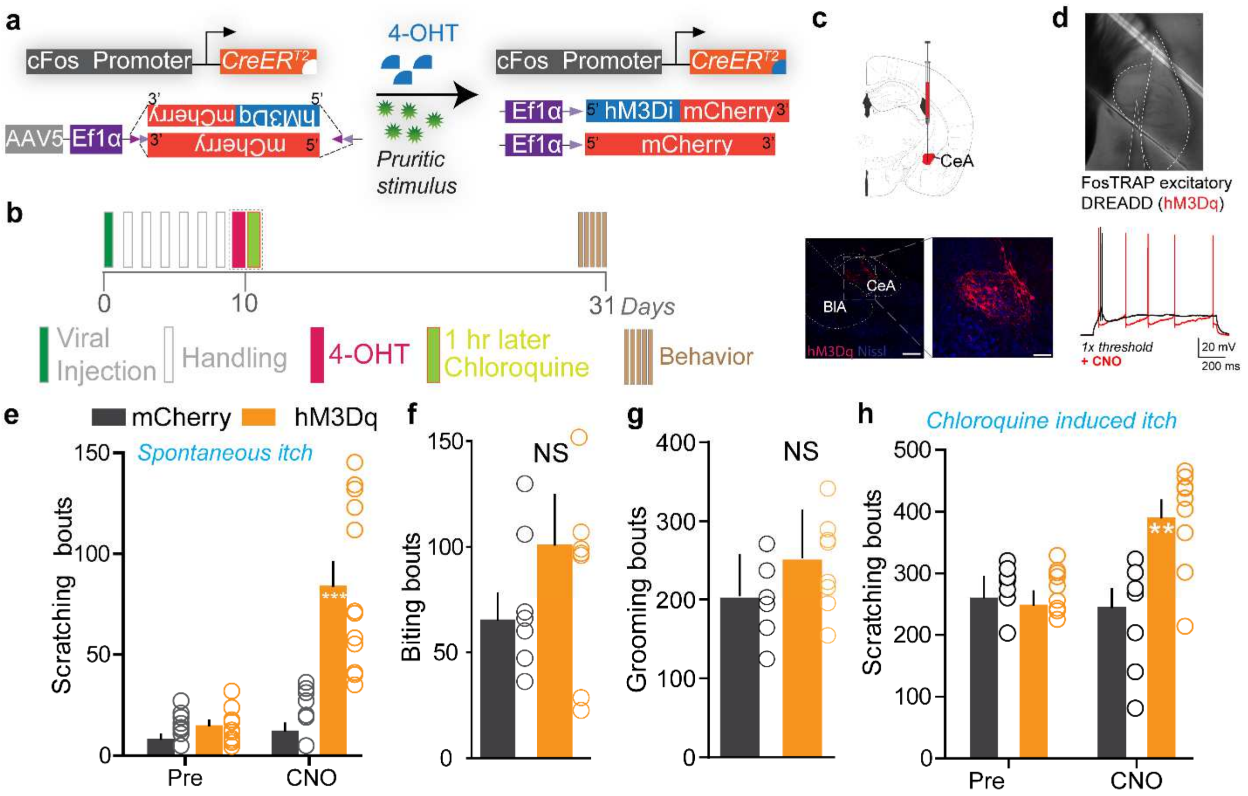


**Supplementary Figure 4. Chemogenetic activation of the CeA neurons promotes itch behaviors.** (a) Viral strategy to selectively express excitatory DREADDs in itch-activated CeA neurons. (b) Experimental timeline to FosTRAP DREADDs in CeA. (c) Representative section showing CeA FosTRAPped neurons expressing hM3Dq-mCherry (Red). Scale bar, 300 µm. (d) IR DIC image of CeA FosTRAPped neurons expressing hM4Di-mCherry. In hM4Di+ve CeA neurons, CNO bath application increased neuronal excitability in response to a 1 second current injection at 1X rheobase. Black trace is pre-CNO, red trace is after bath application of CNO (10 uM) (e) Chemogenetic activation of CeA itch-TRAPed neurons significantly increases spontaneous scratching. CNO had no effect on chloroquine-evoked scratching in mice expressing mCherry. n = 8-11 per group, p=0.0001. Chemogenetic activation of CeA itch-TRAPed neurons had no significant effects on biting (f) or grooming behaviors (g). n=7-8 per group, n-0.0641 for biting, n =7-8 per group, n-0.141 for grooming. (h) Chemogenetic activation of CeA itch-TRAPed neurons potentiates chloroquine-evoked scratching. CNO had no effect on chloroquine-evoked scratching in mice expressing mCherry. n = 9 per group, p=0.0039.

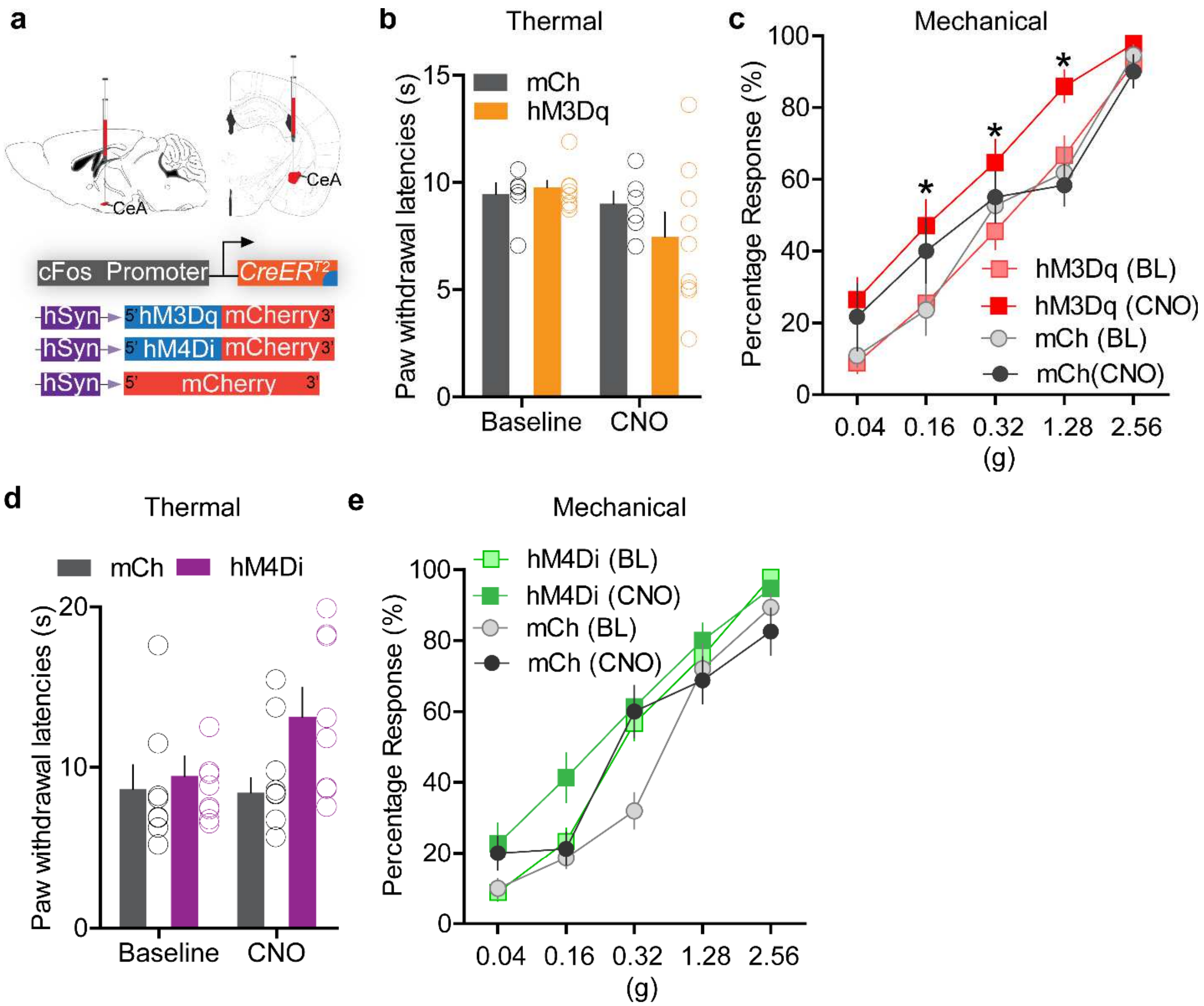


**Supplementary Figure 5. Chemogenetic manipulation of FosTRAPped CeA neurons modulates nociceptive behaviors.** (a) Viral strategy to selectively express excitatory and inhibitory DREADDs in itch activated CeA neurons. (b) Chemogenetic activation of itch-TRAPed CeA neurons does not significantly alter thermal paw withdrawal latencies. n = 7-9 per group, p = 0.46, but significantly increases paw withdrawal sensitivity to mechanical (von Frey) stimulation (c). n = 7-9 per group, *p<0.05. Chemogenetic inhibition of itch-TRAPed CeA neurons has no significant effect on thermal (d) or mechanical (e) sensitivity. n = 7-9 per group, p = 0.22 and p = 0.38.

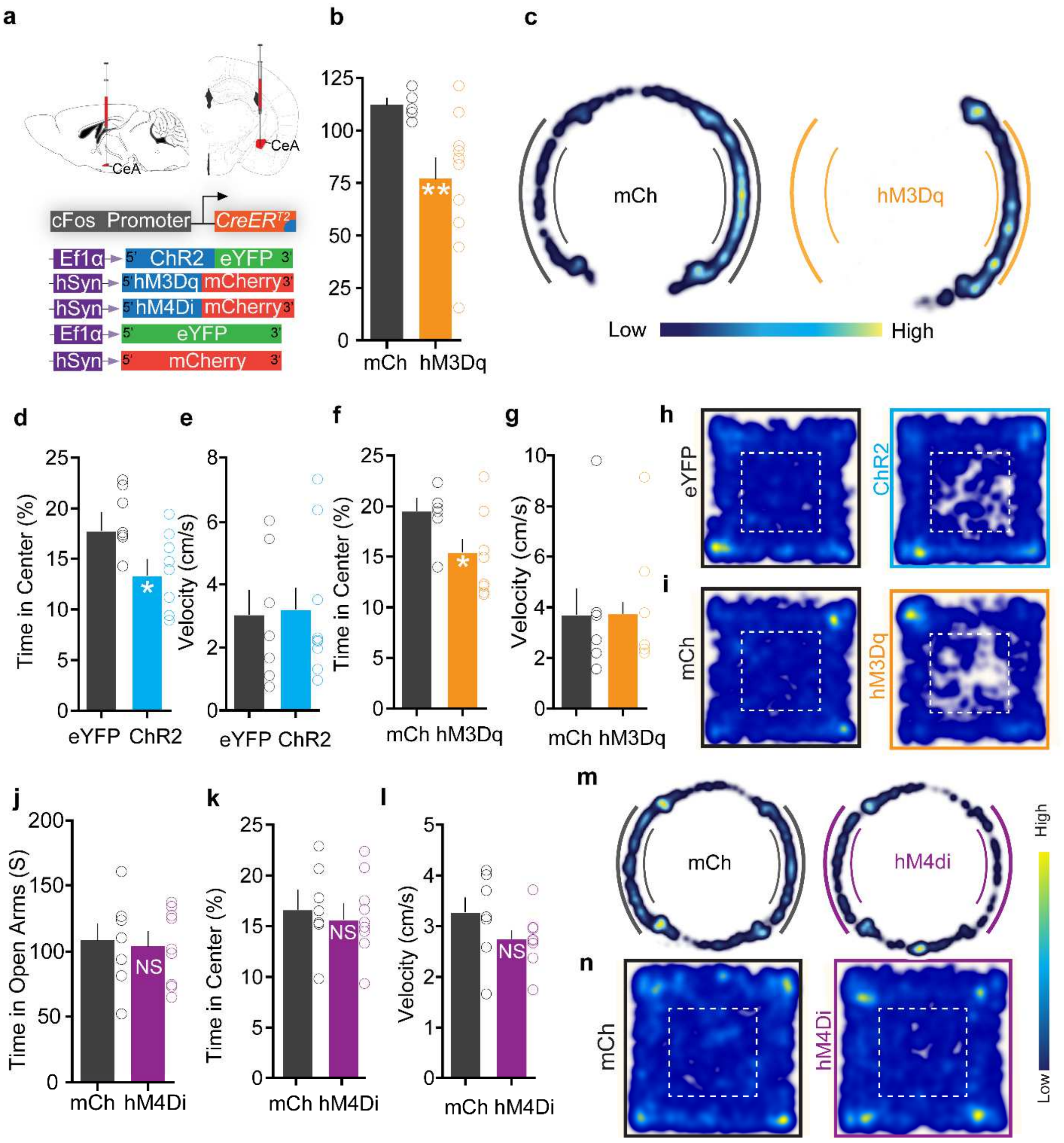


**Supplementary Figure 6. Optogenetic and chemogenetic activation of FosTRAPped CeA neurons causes anxiety but no freezing. Whereas inhibition of these neurons does not affect anxiety state.** (a) Viral strategy to selectively express various constructs in itch-activated CeA neurons. (b) Chemogenetic activation of itch-TRAPed CeA neurons significantly reduces time spent in open arms in an elevated zero maze (EZM). n= 6-10 per group. t test, t=2.439, df=14, p =0.029. (c) Representative occupancy heat map showing spatial location of a control mouse (mCh) and a mouse injected with DIO-hM3Dq in the EZM. (d) Optogenetic and (f) chemogenetic activation of itch-TRAPed CeA neurons causes a significant decrease in time spent in center in the open-field test (OFT). n= 7-10 per group. t test, t=2.86, df=14, p =0.012, for optogenetic group and n= 6-10 per group. t test, t=2.331, df=14, p =0.036 for chemogenetic group. (e, h) Optogenetic and chemogenetic activation of itch-TRAPed CeA neurons had no effect on mean velocity in the OFT. n= 7-10 per group. t test, t=0.4346, df=13, p =0.67, for optogenetic group and n= 6-10 per group. t test, t=1.043, df=14, p =0.31 for chemogenetic group. (h, i) Representative occupancy heat map showing spatial location of a control mouse (eYFP, mCh) and a mouse injected with DIO-ChR2 and DIO-hM3Dq in the OFT. (j) Chemogenetic inhibition of itch-TRAPed CeA neurons has no effect on time spent in open arms in EZM, n= 7-9 per group. t test, t=0.18, df=14, p =0.70; and (k) time spent in center in OFT. n=7-9 per group. t test, t=0.39, df=14, p =0.85 (l) Chemogenetic inhibition of itch-TRAPed CeA neurons has no effect on mean velocity of mice in OFT. n= 8-9 per group. t test, t=1.59, df=15, p =0.13. (m, n) Representative occupancy heat map showing spatial location of a control mouse (mCh) and mouse injected with DIO-hM4Di in the EZM and OFT. n = 8–9 per group.

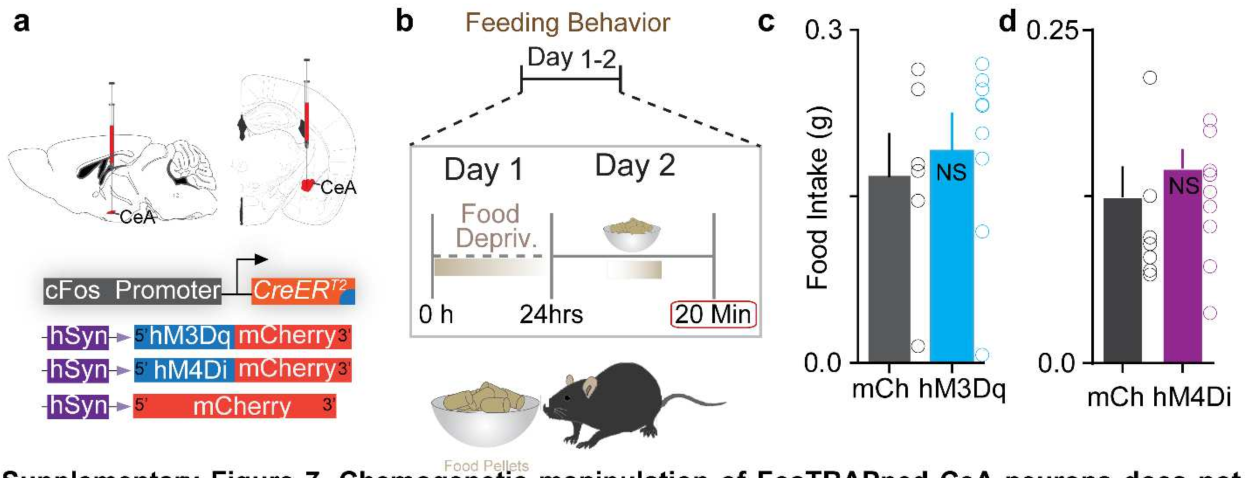


**Supplementary Figure 7. Chemogenetic manipulation of FosTRAPped CeA neurons does not affect feeding behaviors.** (a, b) Viral constructs used and schematic of experimental timeline. Chemogenetic activation (c) or inhibition (d) of itch-TRAPed CeA neurons has no effect on food intake. n= 6-9 per group. t test, t=0.52, df=13, p =0.60 for activation; n= 7-9 per group. t test, t=0.67, df=14, p =0.50 for inhibition.

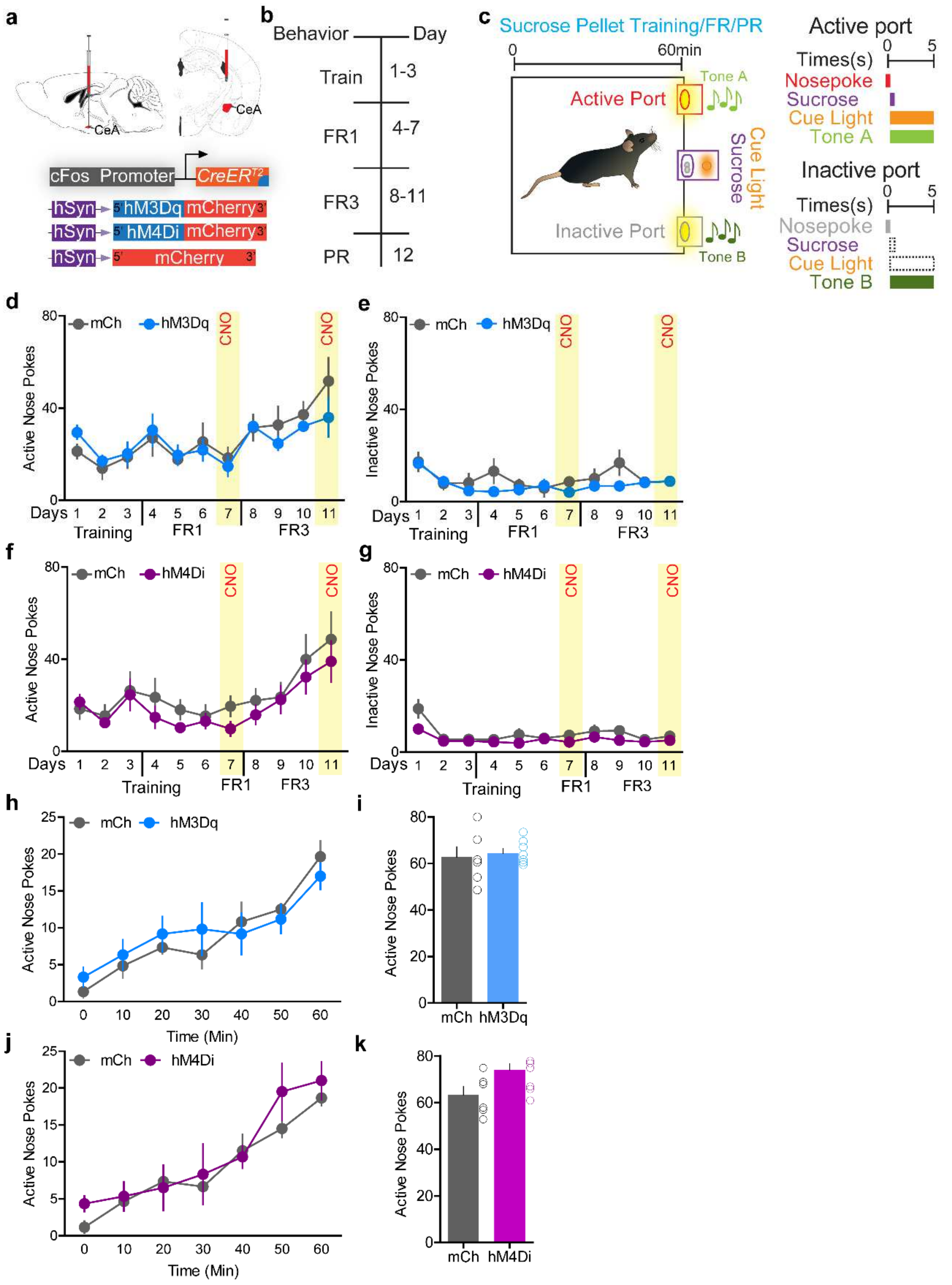


**Supplementary Figure 8. Chemogenetic manipulation of FosTRAPped CeA neurons does not affect reward seeking behaviors.** (a) Viral strategy to selectively express excitatory and inhibitory DREADDs in itch activated CeA neurons. (b) Experimental timeline (FR1 = fixed ratio 1; FR3 = fixed ratio 3; PR = progressive ratio). (c) Sucrose pellet training paradigm. Chemogenetic activation (d, e) and inhibition (f, g) of itch-TRAPed CeA neurons does not alter performance in FR1 or FR3 paradigms where nose pokes are used to obtain sucrose pellets. Activation of these neurons also has no effect on number of inactive nose pokes. For activation (n= 8-9 per group, $F(6,60) = 5.95$, $p = 0.28$ for active nose pokes and $F(6,60) = 1.25$, $p = 0.95$ for inactive nose pokes) and for inhibition (n= 7-9 per group, $F(6,84) = 1.23$, $p = 0.73$ for active nose pokes and $F(6,84) = 8.20$, $p = 0.11$ for inactive nose pokes). (h, I, j, k) Chemogenetic activation/inhibition of itch-TRAPed CeA neurons has no effect on performance in the sucrose progressive ratio (PR) schedule, and also has no effect on number of inactive nose pokes (not shown). t test, $t=1$, $df=10$, $p = 0.62$, t test, $t=1.63$, $df=10$, $p = 0.13$.

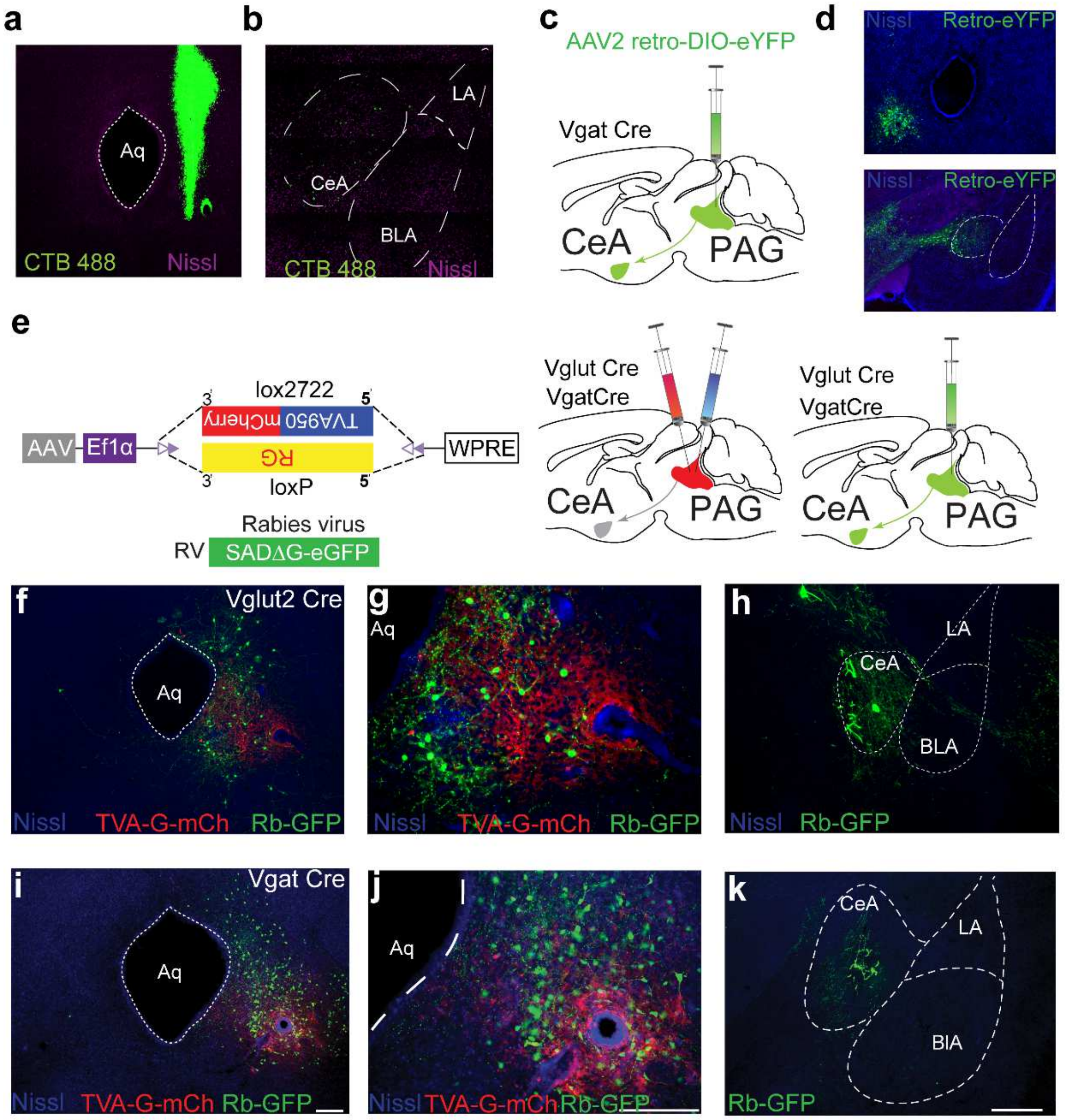


**Supplementary Figure 9. Anatomical tracing to identify connections between the CeA and the PAG.** (a) Retrograde tracer CTB 488 injection into the vPAG to confirm projections from CeA in C57BL/6J mice. Representative image of vPAG showing CTB488 injection. (b) CeA neurons show clear CTB 488 labeling following injection into the vPAG in C57BL/6J mice. (c) Cre-dependent retrograde virus AA2-DIO-eYFP is injected into the vPAG of Vgat Cre mice. (d) CeA neurons show expression of Cre-dependent eYFP in Vgat+ve neurons following injection into vPAG. (e) Strategy for monosynaptic retrograde rabies tracing to confirm the CeA[illegible]PAG connection. (f, g) mCherry fluorescence from Vglut2 neurons selectively transduced with TVA in the vPAG. GFP fluorescence in PAG following SADΔG-GFP(EnvA) injection into the PAG. (h) SADΔG-GFP expression in the CeA following injection into the vPAG in a mouse expressing TVA in Vglut2 neurons. (i, j) mCherry fluorescence from Vgat neurons selectively transduced with TVA in the vPAG. GFP fluorescence in the PAG following SADΔ G-GFP(EnvA) injection into the PAG. Scale bar, 75 µm and 250um. (k) SADΔ G-GFP expression in the CeA following injection into the vPAG in a mouse expressing TVA in Vgat neurons Scale bar, 300µm

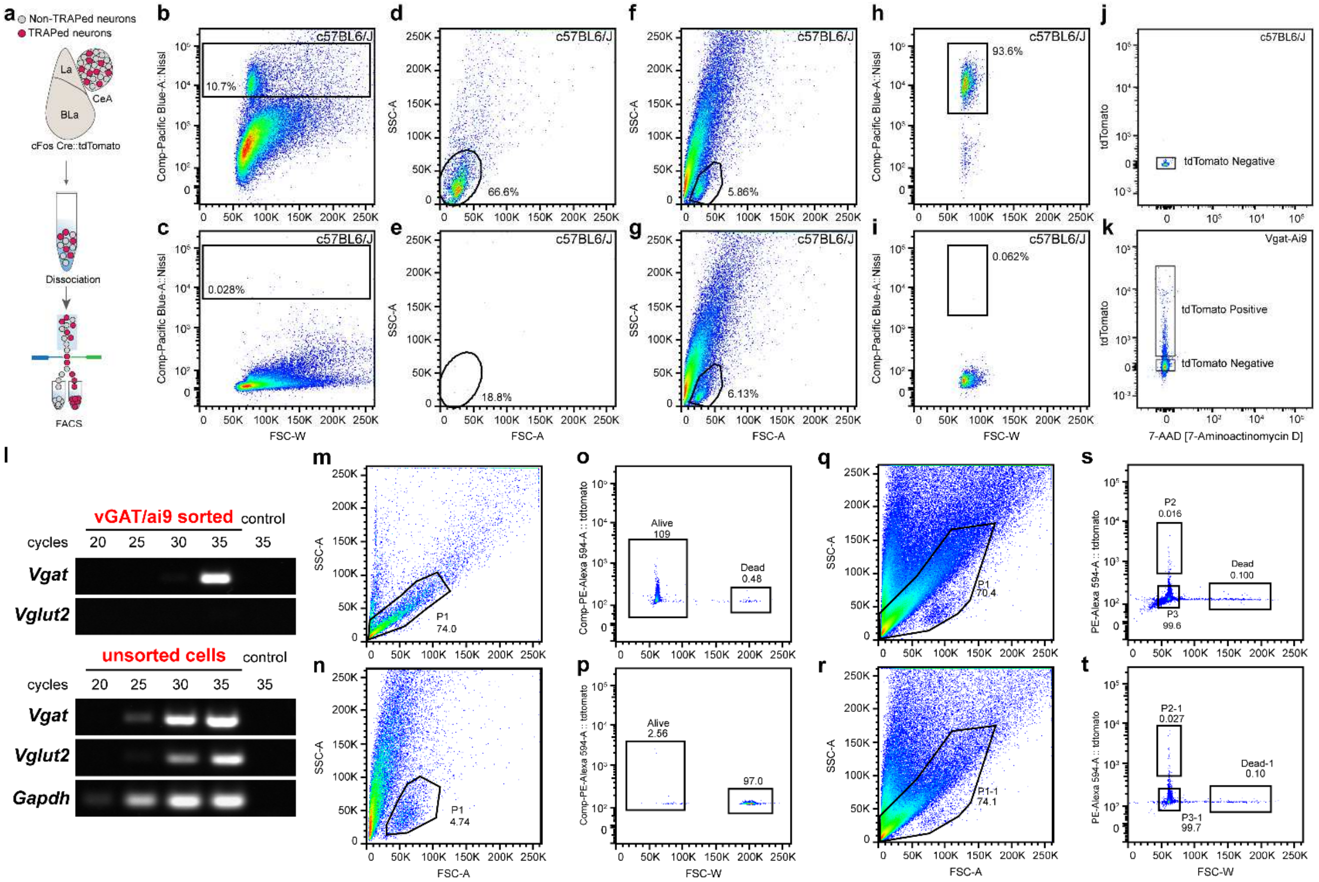

**Supplementary Figure 10. Fluoresence-activated cell sorting of CeA FosTRAPped neurons to perform RNA-seq and transcriptional analysis of itch-TRAPed CeA neurons.** (a) Schematic showing dissociation and FACS of CeA FosTRAPped neurons. (b, d) Labeling of dissociated neurons with Neurotrace reveals a clear subset of events that can be found in the bottom of the scatterplot. (c, e Absence of Neurotrace staining confirms labeling, and rules out an autofluorescence artifact. (f, h) Selection of the identified Neurotrace subset in the FACS scatterplot reveals that it contains 93.6% of all Neurotrace positive events. (g, i Absence of Neurotrace staining confirms labeling, and rules out an autofluorescence artifact. (j) Negative fluorescence FACS control using dissociated tdTomato (Ai9) negative tissue from C57BL/6J animals. (k) Positive fluorescence FACS control using dissociated tdTomato (Ai9) positive tissue from Vgat/Ai9 animals. (l) PCR validation of sorted samples, demonstrating enrichment of population of interest (Vgat). (m, o) Assessment of neuron viability after dissociation. (n, p) Positive dead control for neuronal viability. (q, s) Example of CeA cFos TRAP FACS #1. (r, t) Example of CeA FosTRAP FACS #2.

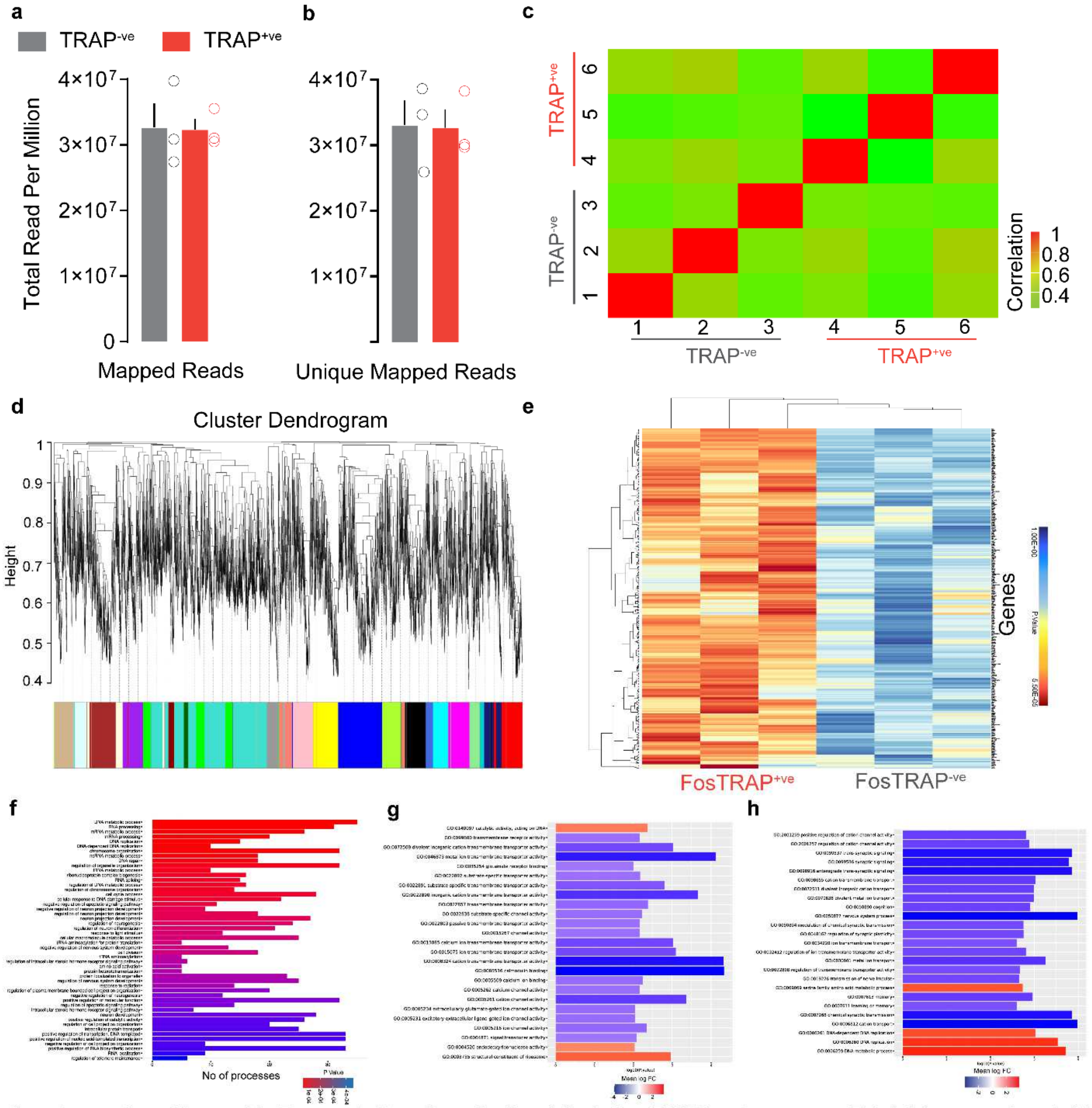


**Supplementary Figure 11. Transcriptional analysis of CeA FosTRAPped neurons.** Total (a) mapped and (b) unique mapped reads per million are similar between tdTomato+ve and tdTomato-ve neurons. t test, t=0.07, df=4, p =0.94 for mapped reads and t=0.08, df=4, p =0.93 for unique mapped reads. (c) Correlation analysis of RNA quality obtained from tdTomato+ve and tdTomato-ve neurons. This matrix supports our prior expectation that the summation of expressed isoforms to the level of their parent genes in the data follows a positive trend with high correlation. This also highlights that there is no cross contamination form different sorting events. (d) Sample cluster dendrogram of tdTomato+ve neurons showing all WGCNA de novo modules of genes identified as random color names where each module eigengene was correlated to tdTomato treatment. The GreenYellow module is uniquely 99% correlated and significant for tdTomato+ve neurons. (e) Heat map of significantly correlated genes filtered from the GreenYellow module. (f) Barplots showing gene ontology biological processes that are significantly upregulated in the tdTomato+ve neurons compared to the tdTomato-ve neurons. (g, h) Top-enriched GO biological processes and molecular function for up-regulated genes (log2FC > 0.5) and down-regulated (log2FC < 0.5) in the tdTomato+ve neurons.
